# Hotspot Dosages of Most Rapid Antibiotic Resistance Evolution

**DOI:** 10.1101/866269

**Authors:** Carlos Reding, Pablo Catalán, Gunther Jansen, Tobias Bergmiller, Phillip Rosenstiel, Hinrich Schulenburg, Ivana Gudelj, Robert Beardmore

## Abstract

We treated *Escherichia coli* with the antibiotic erythromycin from zero to high dosages to determine how the evolutionary dynamics of antibiotic resistant phenotypes and genotypes depend on dose. The most rapid increase in resistance was observed just below erythromycin’s minimal inhibitory concentration (MIC) and genotype-phenotype correlations determined from whole genome sequencing revealed the molecular basis of this: simultaneous selection for copy number variation in 3 resistance mechanisms which shared an ‘inverted-U’ pattern of dose-dependent selection with several insertion sequences and an integron. Many genes did not conform to this pattern, however, because of changes in selection as dose increased: media adaptation at zero-to-low dosages gave way to drug target (ribosomal RNA operon) amplification at mid dosages whereas prophage-mediated drug efflux dominated at higher dosages where population densities were lowest. All dosages saw *E. coli* amplify the efflux operons *acr* and *emrE* at rates that correlated strongly with changes in population density that exhibited an inverted-U geometry too. However, we show by example that inverted-U geometries are not a universal feature of dose-resistance relationships.

## 1 Introduction

To the best of our knowledge, no study has determined which antibiotic dosages promote the most rapid antibiotic resistance adaptation resulting from *de novo* evolution in a population of microbes. We therefore sought the antibiotic dosage of most rapid adaptation in an *in vitro* model by quantifying genomic and phenotypic changes in strains of *Escherichia coli* where the amplification of a genomic region containing the efflux operon, *acr*, provides resistance to the antibiotic erythromycin.^1^ Our experimental procedure was this: for as many antibiotic dosages as is practicable, propagate replicate bacterial lineages at fixed dosages and quantify changes in different fitness measures at each dosage using spectrophotometry, thus quantifying resistance adaptation phenotypically. Then, study the dependence of resistance phenotypes on dose and use longitudinal deep sequence data to correlate resistance phenotypes with genomic change.

Erythromycin is used clinically against Gram negative bacteria, although not to treat *E. coli*. The latter encounters erythromycin as an unintended side-effect of treatment, as do all species in the microbiota. Interestingly, clinical resistance of *Salmonella typhimurium* blood isolates to erythromycin are known to change quickly, including a possible 256-fold increase in resistance from efflux-mediated adaptation within a week^2^ wherein the protein AcrB encoded by *acr* doubled in expression in that time. So while our study is limited by lacking an *in vivo* treatment context, the *E. coli* strains we use are sensitive in the laboratory to erythromycin which can provoke a rapid evolutionary response.

The following idea is key throughout: for any microbial phenotype, *f*, be it population density, exponential growth rate, Malthusian fitness, duration of lag phase, efflux protein expression levels or anything else, *f* will depend on both time, *t*, and erythromycin dosage, *E*. Changes in *f* correspond to changes in antibiotic resistance and so we can use a numerical derivative of *f* to quantify the rate of microbial adaptation.^12^ We can then seek the dosage, *E*, for which this rate is greatest and we call these ‘dosing hotspots’. We can perform analogous calculations using genome sequence data, where *f* is the frequency of some mutation in the population, and so determine the dose-dependence of selection coefficients for that mutation. The purpose of this analysis is to elucidate the ‘geometry’ of the dependence of rates of phenotypic adaptation and selection coefficients for resistance mutations on antibiotic dose.

## 2 Results

We expected the ‘inverted-U’ to appear in the aforementioned geometry, as it has done in prior experiments that compete bacterial strains with different resistance profiles together.^3, 4^ This is the following idea. Selection for resistance increases with antibiotic dose^6^ but antibiotics suppress growth at high dosages. Thus, although mutation rates can increase with antibiotic stresses,^7, 8^ mutational supply could be low enough to forestall resistance at high dosages. If true, this could create a mid-dose regime where resistance adaptation were fastest, whence an inverted U shape would describe the geometry of resistance we describe above where the dosage of most rapid resistance evolution sits at the peak of the U. The inverted U has been described using between-strain competition experiments^4^ but not, so far as we know, using *de novo* mutants.

### 2.1 Theoretical predictions

Theory points to the potential for a greater degree of subtlety in this geometry than just the presence of an inverted U. To see this, consider the following toy genetics model that quantifies the relationship between the rate of resistance adaptation and antibiotic dose:

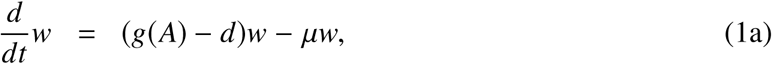

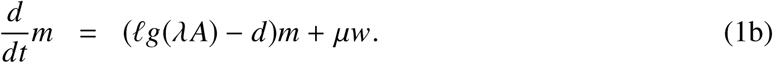

Here *µ* is mutation rate from drug susceptible wild-type bacterial strain (*w*) to resistant mutant (*m*). We assume an initially monoclonal population so that *w*(0) = 1 and *m*(0) = 0, where *g* is *per capita* bacterial growth rate, *d* is a natural death rate, *A* is antibiotic dosage. In terms of dose, *A*, the minimal inhibitory concentration (MIC) of w occurs where *g*(*A*) = *d* and the MIC of *m* occurs where *lg*(λ*A*) = *d*; the MIC of *m* is said to be the mutant prevention concentration (MPC). The scaling parameter λ (between 0 and 1) acts to reduce the drug concentration that *m* experiences relative to the wild-type but this reduction comes at a fitness cost controlled by *l* < 1.

Write *θ*_1_ := *lg*(*λA*) − *d* and *θ*_2_ := *g*(*A*) − *d* − *µ* and note the following exact solution of (1):

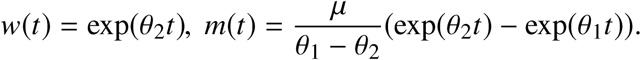

The frequency of antibiotic-susceptible strains in the population, *ρ*, is

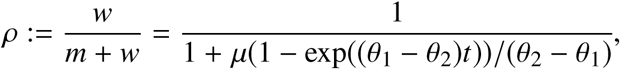

so if *θ*_1_ > *θ*_2_ then *ρ*(*t*) → 0 as *t* → ∞. On the other hand, if *θ*_2_ > *θ*_1_ then mutation-selection equilibrium is reached with *ρ*(*t*) → 1/(1 + *µ*/(*θ*_2_ − *θ*_1_)) as *t* → ∞ and the rate of convergence is determined by

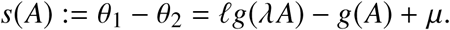

We call *s*(*A*) the *rate of adaptation* (ROA) and the maximum ROA (MRA - *a.k.a.* an ‘evolutionary hotspot’) is the dose, *A*, where *s*(*A*) is maximal. By calculus, the MRA occurs where

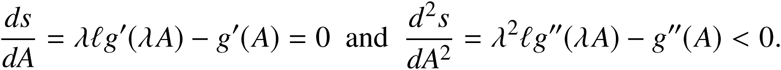

For the case where g is a Hill function, so *g*(*A*) = 1/(*k^p^* + *A^p^*) these equations can be solved but if *g*(*A*) = exp(−*A*)^9^ they cannot, so there is no inverted U in this case. In the former case where an inverted U is present, so too a MRA, then this calculation shows the location of the MRA depends on λ and *l*, predicting the MRA dosage depends on growth rate costs of resistance and the reduction in antibiotic dosage found inside a cell due to a resistance mutation. As different mutations are likely to have different costs of resistance, this model predicts, therefore, that different mutations will have different MRA hotspots.

A more sophisticated density and frequency-dependent population genetics model^10^ tuned to erythromycin efflux makes a prediction we can seek in data: redolent of the Eagle Effect,^11^ in drug-adapted dose responses, where population density is regressed against antibiotic dose following each treatment, do not remain monotone as they increase in density to form a ‘bump’ at the MRA hotspot. That model predicts short-term theoretical dose-responses follow standard antibiotic sensitivity assays whereby the addition of more drug reduces bacterial density,^10^ but repeated antibiotic treatments could modify the dose response such that it appears non-monotone at a hotspot due to selection for subpopulations with amplified pump operons in their genomes.

### 2.2 Patterns of selection in data: summary

To test these theoretical ideas, we determined the geometries of selection for resistance by treating 4 strains of *E. coli* K12 (AG100, AG100A, TB108 and eTB108) with antibiotics, deploying erythromycin at several concentrations (0, 5, 10, …., 40µg/mL) daily for 7 days (*a.k.a.* seasons, 8 replicates, see Methods) and measuring population densities and efflux protein expression continually (TB108 and eTB108 only, as explained below). We then quantified evolved dose responses for all strains and sought the resistance mechanisms supporting them by sequencing 3 AG100 metapopulations sampled from 3 replicate lineages every other day.

To summarise our findings, AG100 exhibited dose-dependent evolution where the accumulation of *de novo* single nucleotide polymorphisms (SNPs) in genes with different functions decreased in number as antibiotic dosage increased. Amplifications of the drug targets (7 rRNA *rrl* operons) and 2 efflux operons (*acr* and *emrE*, the latter encoded within a prophage) increased in frequency at different rates at different dosages, as did several *ins* (insertion) sequences and the integron *intD*. Different rates of sweep of resistance mechanisms created different inverted-U geometries: *rrl* amplifications were selected most near 15 − 20*µg*/*ml* whereas amplification of the efflux operons were selected most near to the MIC, namely 25 − 30*µg*/*ml*. Efflux mechanisms were amplified at all positive antibiotic concentrations, including the lowest, and mutants with at least 3 copies of the *acr* operon swept to fixation within 72h, a process that happened fastest close to the MIC.

The change in population density data of AG100 exhibited an inverted-U geometry whereas strain AG100A lacks a functional AcrAB-TolC efflux pump and it did not exhibit an inverted-U, thus the inverted-U is not a universal feature of adaptation to an antibiotic. Strains TB108 and eTB108 are derived from *E.coli* MG1655 and have GFP phsyically fused to AcrB, they therefore provide approximate data on AcrB expressed per cell at all times. TB108 data show that hotspots in population density adaptation correlate with changes in efflux pump protein (AcrB) expression (*R*^2^ ≈ 0.84) that correlates positively with *acr* operon amplification; thus *acr* explains the lion’s share of the resistance adaptation.

### 2.3 Patterns of selection in detail: the MRA can be close to the MIC

We measured optical densities of AG100 populations continually and they formed a standard dose response that declined with increasing drug concentration (Figure 1A, 24h data), putting the MIC (the IC_99_) at 32.1 ± 1.8µg/mL (mean ± 95% C.I., *n* = 8; Figure S1). By season 3, AG100 was detected at dosages where it previously had not been (Figure 1A, 72h) and all populations exhibited increasing densities, indicating reduced antibiotic efficacy (Figure 1B).

**Figure 1.**
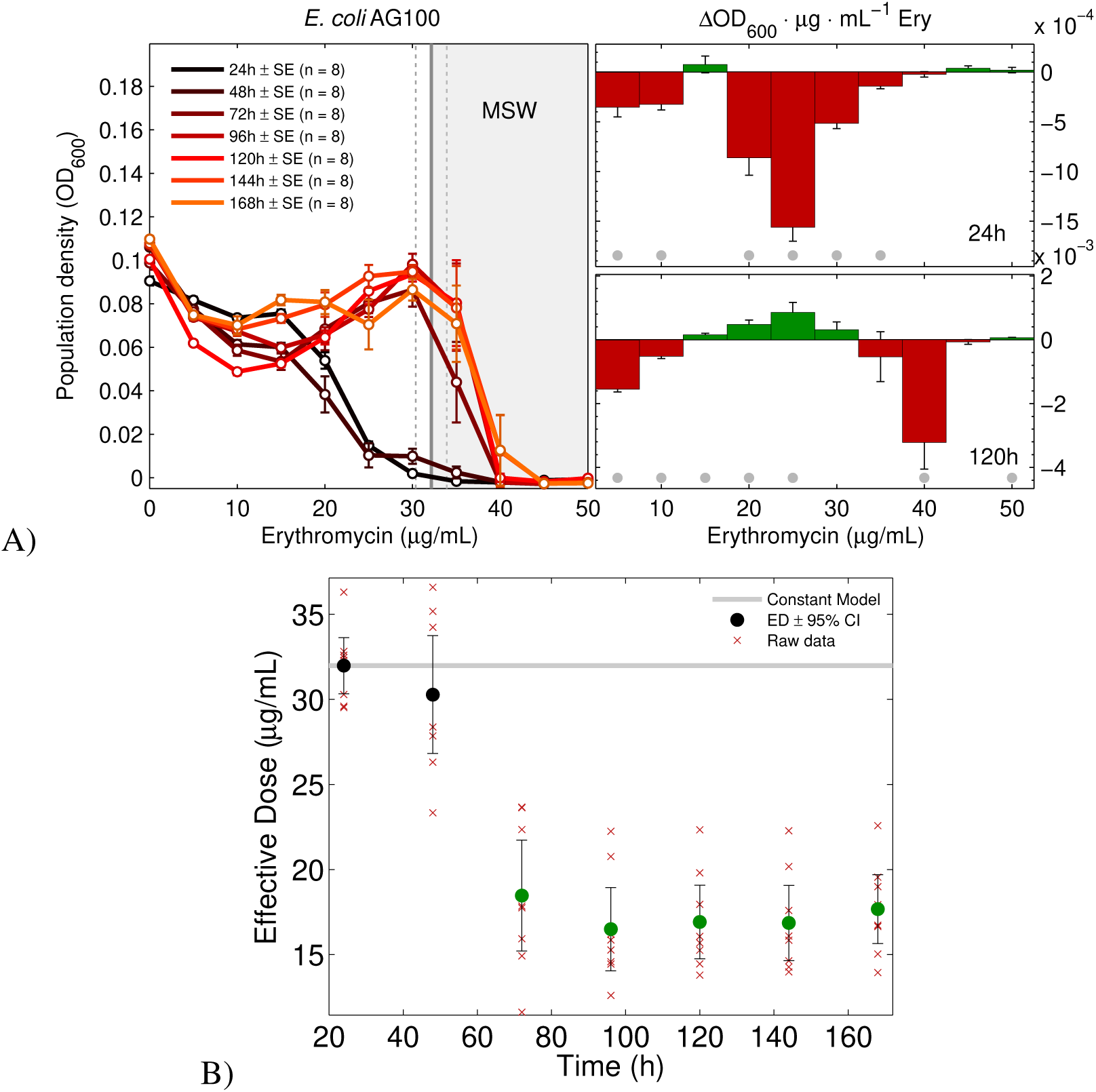
Evolving a dose response using AG100. **A)** Evolving *E.coli* AG100 dose-response profiles plotted each 24h season. The MIC is the vertical line (32.1*µg*/*ml*± 95% CI (dotted vertical lines)), the hypothetical MSW is the grey area. Increments in drug concentration can increase (green bar) or decrease (red bar) bacterial density, as indicated in the right subplot after 24h and 120h of exposure to erythromycin (mean ± standard error, n = 8; significant changes (*p* < 0.05) with respect to *H*_0_*: no change induced by increasing drug concentration based on a two-sided t-test* are denoted by a grey dot. **B)** The reduction in effective antibiotic dose (defined in Methods) for AG100. The null *H*_0_*: sensitivity to erythromycin does not change through time* is the grey line; green dots are indicative of significant changes beyond 72h using a 2-sided t-test (*p* < 0.05).

To quantify rates of adaptation (ROA), we applied a rate of change measure to population densities and growth rates.^12^ Three versions of this measure (Methods) concur that adaptation to erythromycin has a non-linear, non-monotone dependence on dose, consistent with prior theory,^10^ and populations adapted fastest when exposed to near-MIC dosages (30 and 35*µg*/*ml*, Figure 2A); the latter figure reveals our first inverted-U. eTB108 behaves analogously (Figure 2) with a strong correlation between ROA in population density (Figure 2A) and ROA of population mean GFP per optical density (Figure 2B; Deming regression *R*^2^ ≈ 0.84, p « 0.05, Figure S2), so increases in AcrB expression correlate with population density.

**Figure 2.**
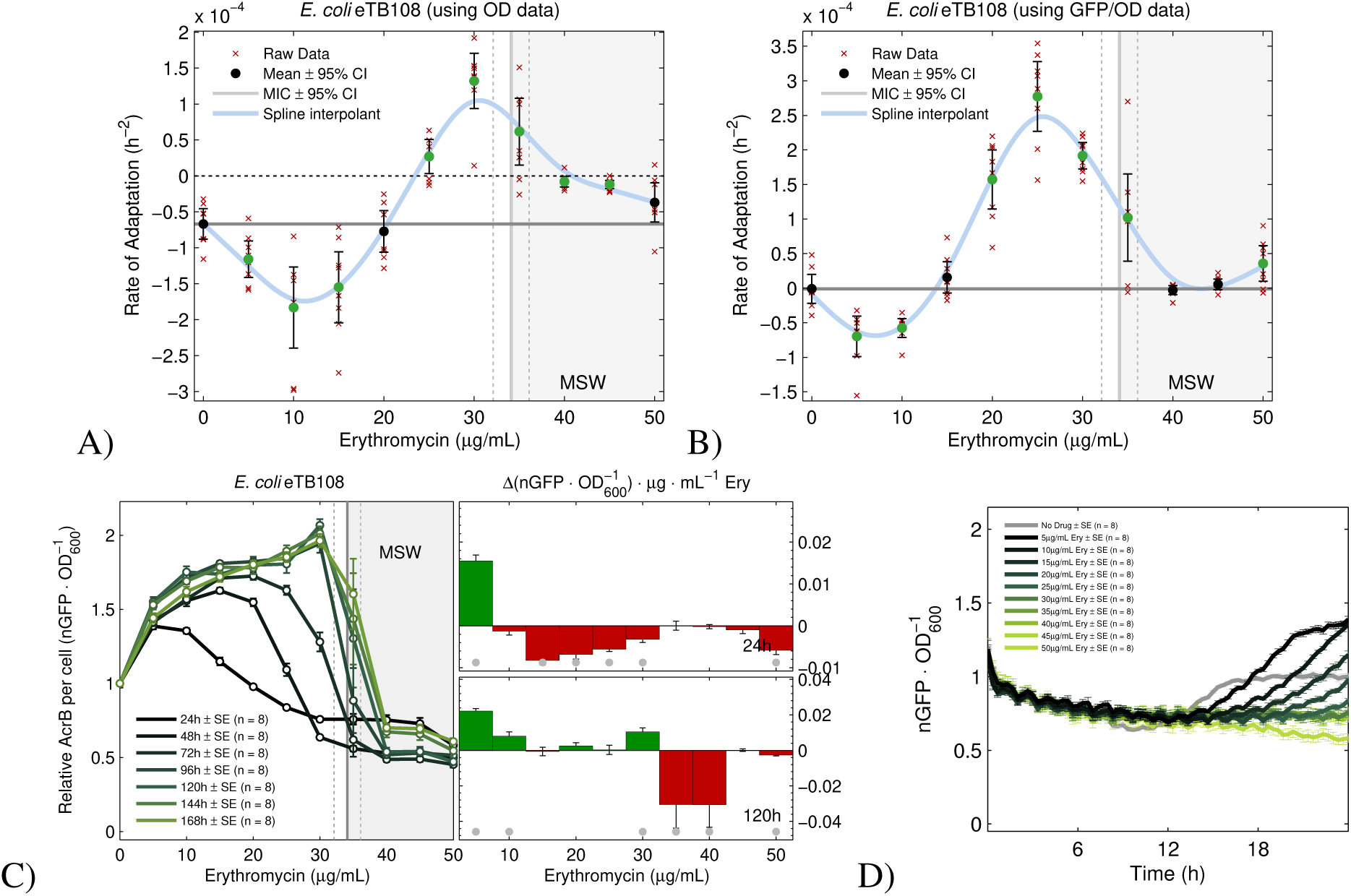
Rates of adaptation depend on drug concentration. **A)** Rates of population density adaptation of eTB108 vary with drug concentrations, notice how a quadratic spline interpolant of the data mean peaks near the MIC (Figure S20 shows analogous data for AG100). **B**) Rates of adaptation of population mean AcrB-GFP expression correlates with population density adaptation (*R*^2^ ≈ 0.84, Figure S2). A quadratic spline interpolant of the data mean is blue, dashed horizontal lines indicate zero adaptation, thick horizontal lines indicate adaptation rate in the absence of erythromycin. Significant changes with respect to the latter are denoted by green dots (two-sided t-tests, mean ± standard error, n = 8). **C)** Green lines are season-by-season experimental dose-response profiles of population mean AcrB-GFP per cell. The effect drug concentration increments have on AcrB-GFP per cell is shown in the right subplots at 24h and 120h of exposure to erythromycin (mean ± s.e., n = 8): increases are green bars and decreases are red bars. Significant changes, with respect to *H*_0_*: no change in mean AcrB-GFP accrues when increasing drug concentration*, based on a two-sided t-test are denoted by a grey dot: note the change from predominantly negative to positive correlations between drug dosage and mean AcrB expression. **D)** Mean AcrB-GFP expression per cell measured for one 24h season using eTB108 exposed to different erythromycin (Ery) concentrations for 24h (mean ± standard error, n = 8; oscillations seen in data are a mechanical phenomenon (Figure S19)).

So as to elucidate increases in resistance of each adapted lineage, one could sample these each day and perform dilution-based assays to determine their respective MICs. However, instead, we introduce an ‘effective antibiotic dose’ measure (EAD, Figure 1B, see Methods) which is the dose that, if it had been applied to the un-evolved wild-type, would have resulted in the same population density as the drug-adapted population; this has the advantage over MIC assays that it can be determined directly from data. Now, for the fastest adapting populations treated at MIC dosages, the EAD reduces by approximately 50% within 72h (Figure 1B).

We re-implemented the 7-day treatment protocol using *acr* knockout strain AG100A and this does not exhibit an inverted U: neither the dose response monotonicity nor the EAD changed significantly (Figure S3B and S4) during treatment.

#### Changing correlations between antibiotic dose and AcrB expression

Given the importance of *acr*, we now address how the expression of protein AcrB adapts during treatment. Seeking relationships between drug dose and GFP tagged AcrB expression, we used eTB108 to quantify changes in mean AcrB-GFP expression per cell at different erythromycin dosages. As erythromycin binds to the ribosomal 50S subunit, inhibiting protein synthesis,^13^ this inhibitory property might be sufficient to create a negative correlation between dosage and AcrB-GFP expressed. However, this correlation is, in practise, not negative but nuanced and changeable. For instance, mean AcrB-GFP per cell at 24h increases significantly as drug passes from 0 to 10*µg*/*ml* (Figure 2C) but AcrB-GFP levels decrease as drug is increased further, up to and beyond the MIC; this remains true in late phase bacterial growth (12-18h, Figure 2D). However, this pre-24h, predominantly negative pump-drug correlation is replaced in a step-wise manner by an increasingly positive correlation as treatment proceeds, creating a wave-like front in changing AcrB expression profiles (Figures 2C, S5).

#### *acr* amplification correlates with loss of drug efficacy and increased AcrB expression

AcrB expression tends to increase each treatment round (Figure S6) although AcrB-GFP declines each day during the first hours after treatment begins (in lag phase) only to increase later through logistic-like dynamics, and the latter occurs whether erythromycin is present, or not (Figures 2D, S6). AcrB-GFP expression dynamics are similar in the absence and presence of antibiotic (Figure 2D), indeed *acr* is known to help bacteria negotiate different phases of growth.^14^

Changes in *acr* copy number per genome have a predictable effect on AcrB expression. To show this we used whole genome sequence (WGS) data to correlate chromosomal copies of *acr* with AcrB-GFP data, we also compared rates of adaptation of population density with *acr* copy number inferred from DNA coverage as a proxy (Methods). Accordingly, 5 days’ erythromycin exposure selects for an amplified 302kb chromosomal region^15^ (274–576kbp) containing *acr* (480– 485kbp) whereby just one copy of this region is observed at all times in the absence of antibiotic (Figures 3, S7–S9). When drug was present, *acr* copy number increased at rates that depended on dosage (Figure 4A and 4B) where the highest rate occurred at around 30 µg/mL (Figure 4B). For comparison, Figure 4C shows amplifications of the ribosomal RNA operon *rrlB* with its lower rate of sweep.

**Figure 3.**
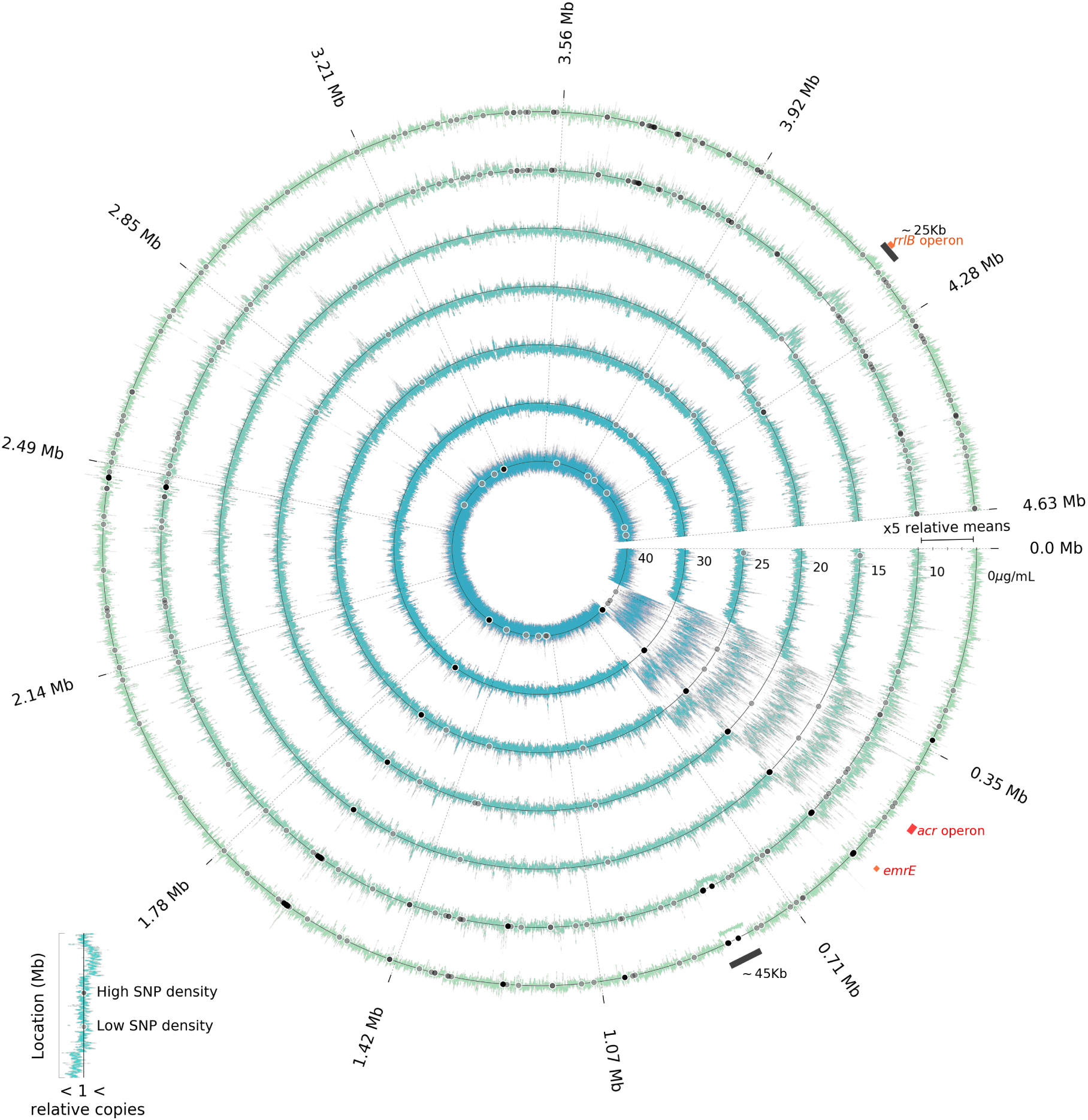
Whole genome sequencing data of AG100 at different erythromycin concentrations following 5 days’ exposure. *Copy number variation*: normalised genome coverage data of populations exposed to different erythromycin concentrations: the outermost ring is for populations grown in the absence of antibiotic, subsequent inner rings show data for increasing concentrations of antibiotic (10-40µg/mL; for replicates see Figures S10 and S11). Coverage data in excess of the mean indicates gene amplification, values below the mean indicate gene deletions, novel single nucleotide polymorphisms above 5% frequency are indicated by black dots indicating regions of evolutionary parallelism and divergence between treatments. The amplified operon *acr* encoding AcrAB is highlighted in red and the amplification of a ∼25kb region between 4,164–4,189kbp is highlighted in black, this region contains metabolic genes, the ribosomal operon *rrlB*, the 50S ribosomal protein L1 (*rplA*) and the RNA polymerase subunit β.

**Figure 4.**
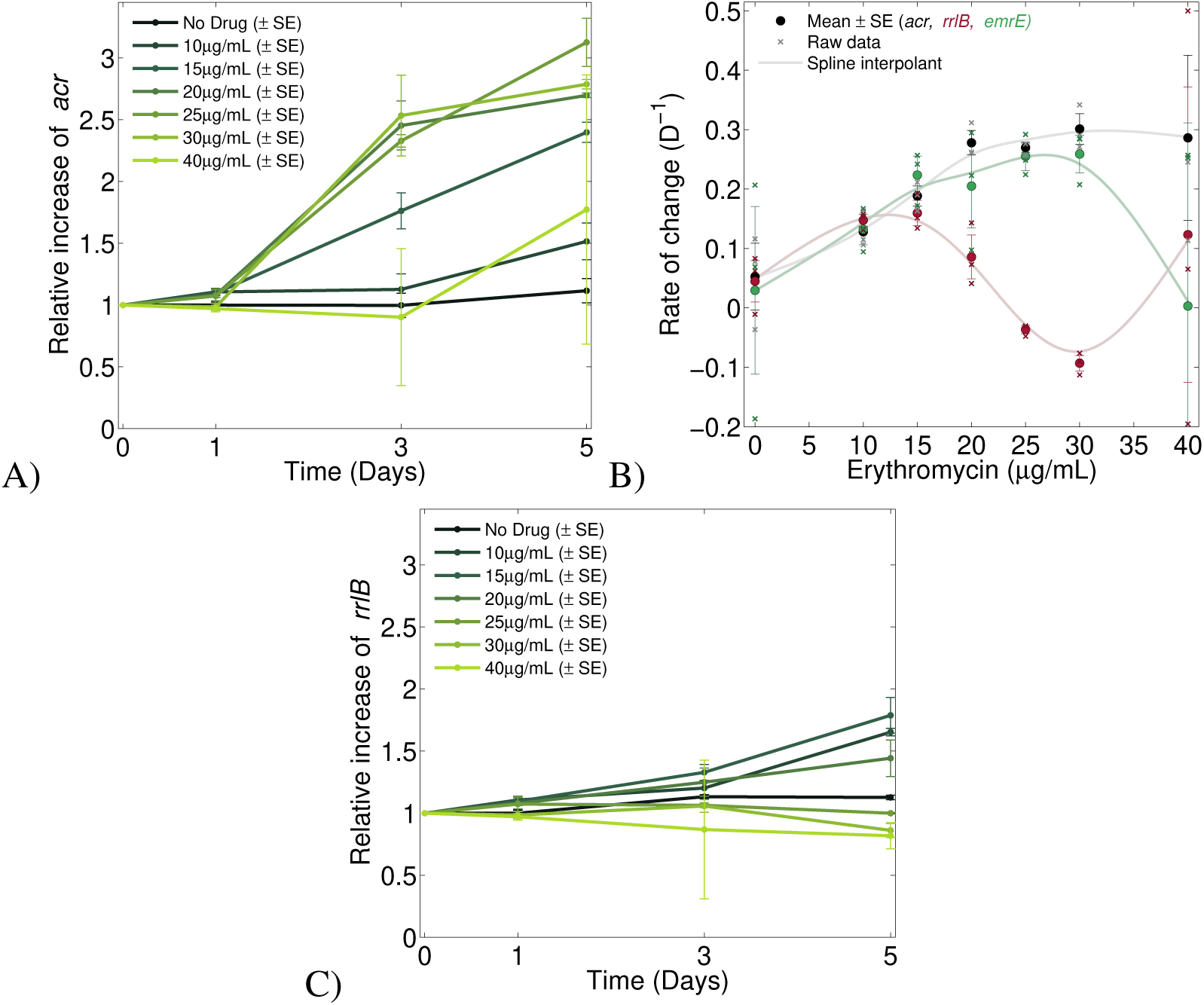
3 inverted-U geometries: selection for *acr, emrE* and *rrlB* amplifications are maximised at different dosages. **A**) Relative abundance of *acr* per genome through time (mean ± s.e., n = 3) at different erythromycin concentrations (colour coded, see legend). **B)** Rates of increase of *acr, emrE* and *rrlB* copy number at different antibiotic concentrations: these are maximised at 30*µg*/*ml* for *acr* and *emrE* and at 15*µg*/*ml* for *rrlB*. **C**) The analogous plot to A for the drug target *rrlB* indicating positive selection for *rrlB* duplications.

*acr* amplifications correlated with increases in AcrB-GFP expression (AcrB per cell: Figure 5A, AcrB per operon: Figure 5B) with significantly positive, nonlinear correlations (AcrB per cell versus operon copies: Figure 5C). As erythromycin inhibits translation, we reasoned the number of efflux pumps per cell per *acr* operon should decline with increasing antibiotic dose and data concur (Figure 5B). We then reasoned both *efflux pumps per cell* and *pumps per cell per operon* (regulatory adaptation) could increase during adaptation? However, no mutations were observed in known regulators of *acr*, including AcrR and MarR. However, we did observe the following.

**Figure 5.**
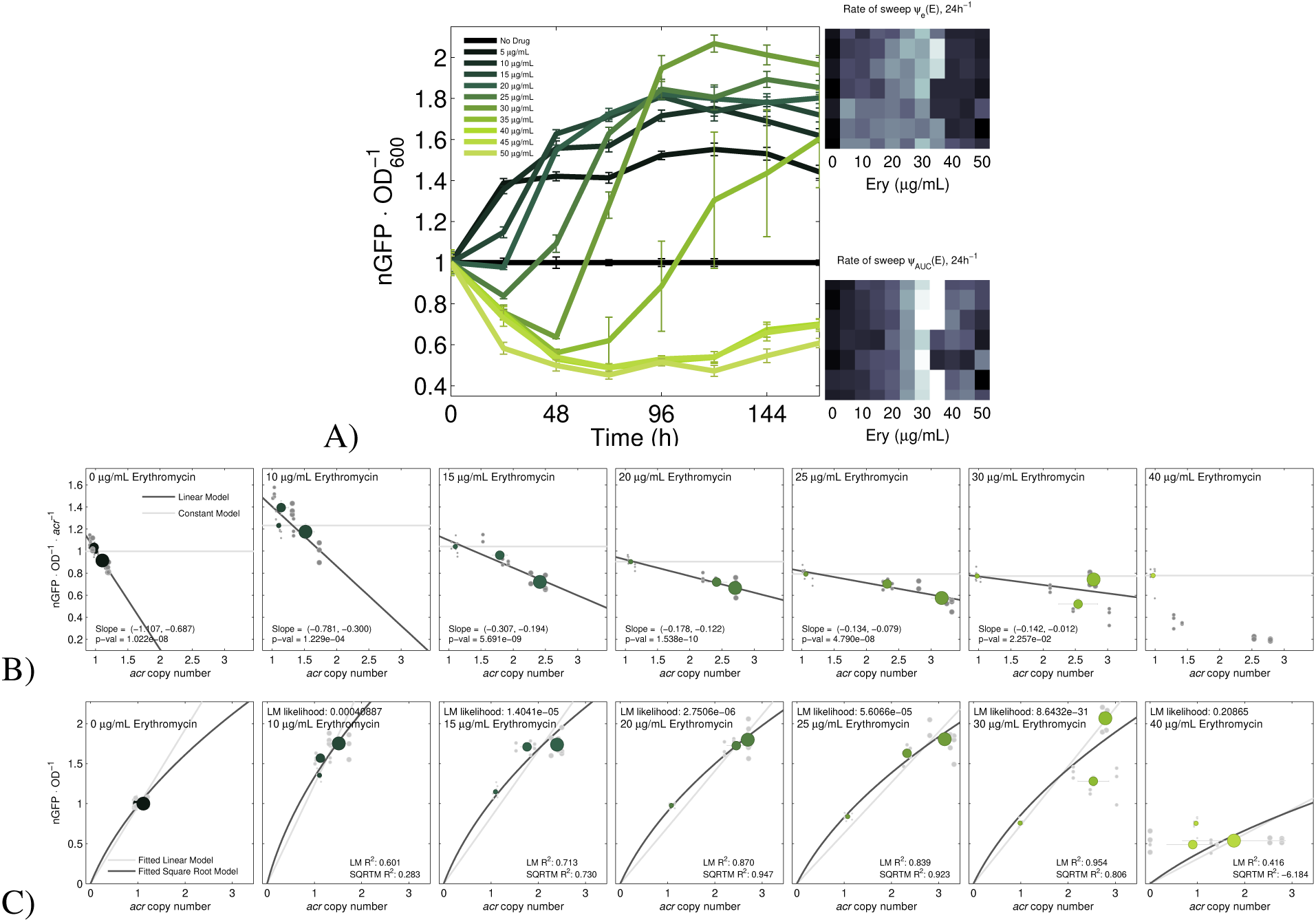
Increasing *acr* copies in the genome increases AcrB expression. **A)** Temporal dynamics of population mean AcrB-GFP per cell at different concentrations of erythromycin (colour coded, mean ± s.e., n = 8). Rates of increase in AcrB-GFP levels are shown in the rightmost greyscale heatmaps using two metrics (see Methods) and are maximised at near-MIC concentrations (for each of 8 replicates, a greyscale image uses a white block for fast adaptation and black for slow). **B)** As erythromycin inhibits translation, population mean AcrB-GFP per cell per *acr* operon decreases with increasing drug concentration (Figure 2C(right) provides statistics, Figure 2D shows these effects in different phases of growth). However, adaptation to antibiotic ameliorates that decrease, as shown by linear regressions whose negative slopes approach zero from below during periods of adaptation to drug. Horizontal grey lines indicate AcrB-GFP produced per operon in the absence of drug, raw data are grey dots, means and s.e. (n = 3) for each day are green dots of increasing size (where size indicates 24h, 72h and 120h). **C)** AcrB-GFP per cell (y-axis) exhibits a positive correlation with *acr* per genome (x-axis) in different erythromycin backgrounds ranging from no drug on the left-most plot (where no correlation is reported due to an absence of *acr* duplications) to super-MIC on the rightmost plot.

In the absence of erythromycin, little daily variability in *acr* per chromosome and AcrB per cell was observed, both maintained wild-type levels throughout treatment (Figures 4A and 5A). However, both varied when drug was applied: at 5-10µg/mL of antibiotic, mean AcrB-GFP per cell increased and reached an apparent equilibrium level approximately 50% above wild-type levels (Figure 5A). Mean AcrB-GFP per cell increased further at higher drug concentrations, achieving double wild-type levels (Figure 5A). The rate of increase of AcrB per cell depended on antibiotic and AcrB per cell reached its equilibrium level fastest at dosages close to the MIC (30*µg*/*ml* in Figure 5A). Moreover, the doubling of AcrB expression was associated with populations in which *acr* per chromosome had tripled (Figure 5C). This is consistent with the auto-repression of *acr* by AcrR whereby fewer pumps per cell are produced per additional operon (linear regressions indicate negative correlations for this, Figure 5B). However drug exposure mitigates this to the point where repression of *acr* is all but lost at near-MIC dosages within 5 days (i.e. regressions showing pumps per *acr* operon (Figure 5B) have negative slopes that increase towards zero as antibiotic dose increases). Despite this phenotypic evidence of regulatory adaptation, no mutations in known regulators of *acr* were found.

#### Genomic amplifications under selection

Coverage data sampled at 24h, 72h and 120h reveals several genomic regions whose amplifications increase in frequency during treatment. The rate of sweep of amplifications of the *dlp12* prophage-encoded efflux pump, *emrE*, exhibits an inverted-U when regressed against antibiotic dose in a manner analogous to *acr* (Figure 4B). Operon *rrlB* experiences antibiotic-dependent amplification with an inverted-U (Figure 4B and C) but the MRA for all *rrl* operons occurs near the sub-MIC dose of 15*µg*/*ml* erythromycin (Figure 6A). Figure 6 summarises selection coefficient data for the amplification of all *rrl* operons, *acr* genes, *emrE*, all *ins* insertion sequences and all *int* integrons. Where there are multiple copies of any of these in AG100, selection coefficients are summed across all copies which is denoted in Figure 6 by a sigma, Σ. Figure 6 highlights genes whose copy number changes have inverted-U geometries, all with dosages of fastest increase from 15 to 30*µg*/*ml*.

**Figure 6.**
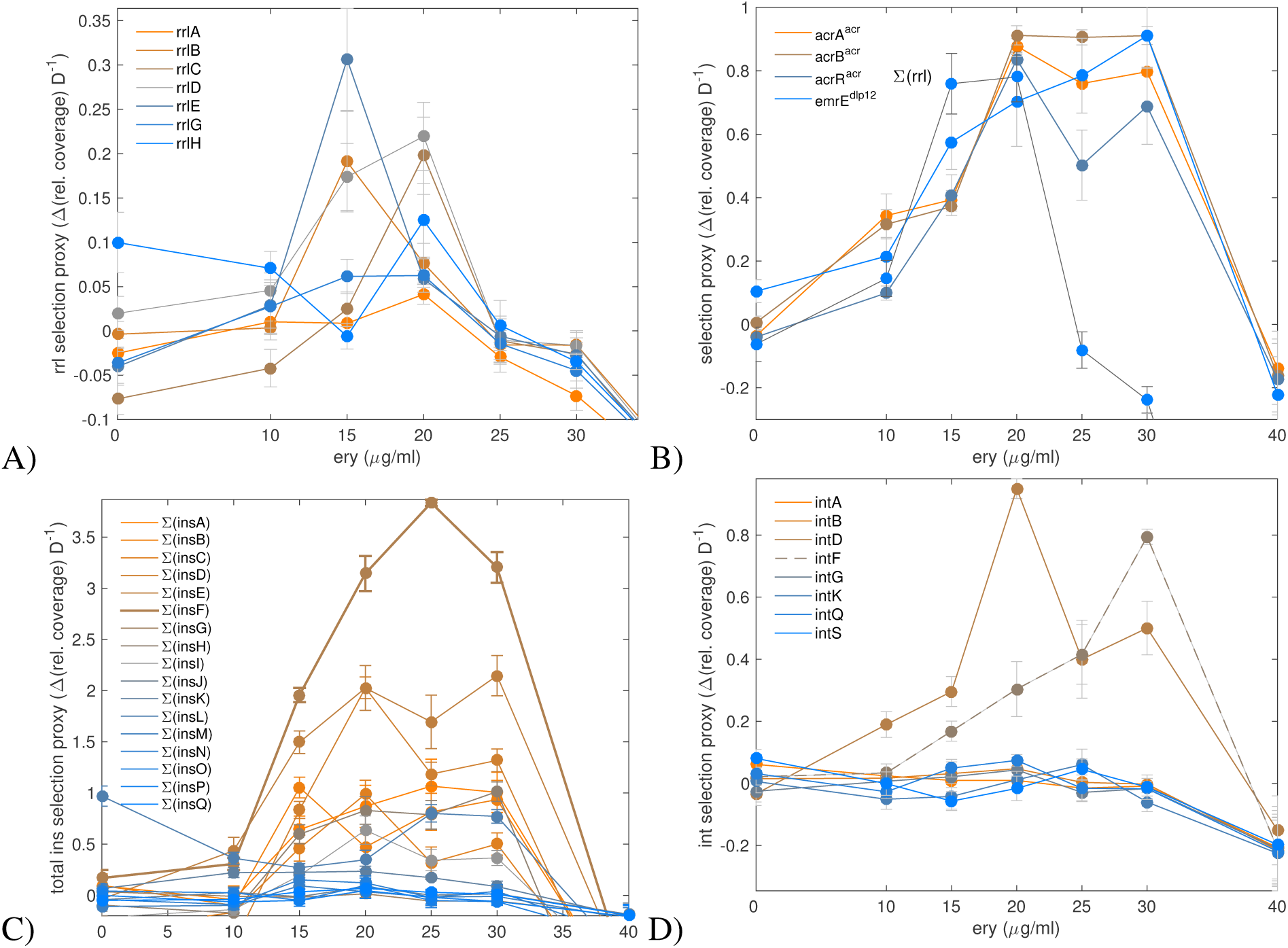
Genes where selection for increased copy number has an inverted-U geometry. A) As shown for rrlB in (Figure 4B) selection for increased copy number of the rrl operons during erythromycin treatment peaks around 15*µg*/*ml*, which is below the MIC at around 30*µg*/*ml*. B) The analogous raw selection profile data for the *acr* operon and *emrE* gene. C) Raw selection profiles for insertion sequences where the selection coefficient data have been summed across all copies of each sequence found in the wild-type genome. Not all IS sequences are amplified in the genome, indeed most are not. D) The integron *intD* has an inverted-U selection profile, as does *intF* (dashed line) but the latter is sited close to *acr* in the genome and has a selection profile (see B) analogous to that operon.

However, Figure 6 also shows genes that do not exhibit an inverted-U pattern. To investigate this variability in more detail, we sought the antibiotic dosages at which the amplification of every *E.coli* gene was maximal. From this we found (Figure 7A) clusters of genes containing the operons *rrl* (MRA around 10-15*µg*/*ml*) and *acr* (MRA close to 20*µg*/*ml*) and the *dlp12* prophage (MRA close to 20*µg*/*ml*). Amplifications of the Qin prophage, however, have no inverted-U and instead see selection coefficients decline with antibiotic dose (Figure 7B) and so Qin may be implicated in adaptation to media, not to the antibiotic.

**Figure 7.**
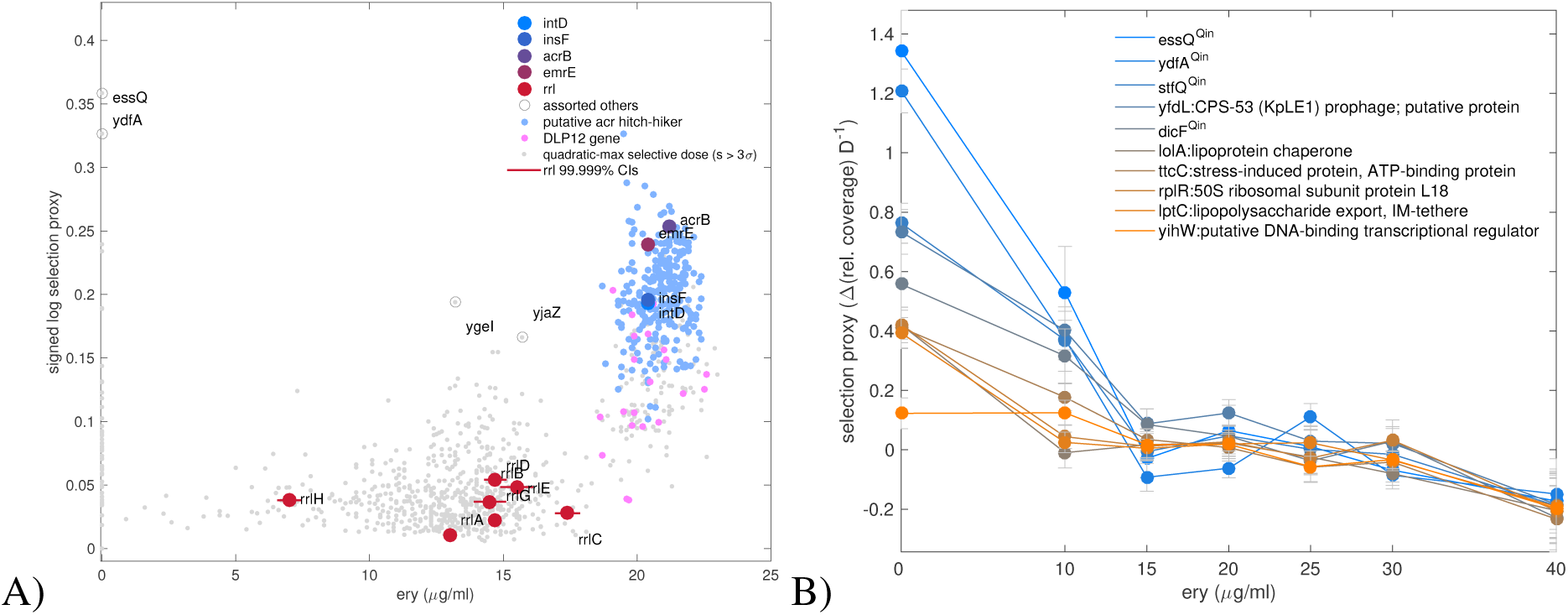
Selection for the amplifications of different genes are maximised at different erythromycin dosages. A) Let *s*(*E*) be the selection coefficient of the genomic amplification of an *E.coli* gene, where *E* indicates the dependence of this quantity on erythromycin. Shown are predicted erythromycin dosages of maximal selection, *s* = max*_E_*_=0…40*µg*/*mL*_ *s*(*E*) (with *E* on the x-axis) for genomic amplifications of every *E.coli* gene where *s* is significantly positive (one datum for each gene observed 3 standard errors above 0 in a quadratic regression, see Methods). Two main clusters are apparent: one (blue and pink dots) shows selection is maximal near the MIC for genes associated with the amplification of regions containing *acr* and *emrE* efflux pumps, along with other genes sited within the *dlp12* prophage, insertion sequences (*ins*), integrons (*int*) and a large, scaleable genomic region that contains *acr*. This cluster lies just below the MIC dosage whereas another containing the *rrl* operons (grey and purple dots) shows maximal selection for the amplification of these occurs at lower dosages. Moreover, the selection profiles of the *rrl*, *ins* and *int* data differ (detailed in Figure 6A). Dosages of maximal selection for copy number variation in other genes reside at both mid and low dosages, including *essQ* from the Qin prophage that exhibits maximal selection at zero erythromycin dosage (other genes in B consistently show selection on Qin amplifications decreases as erythromycin dosage increases). B) Shown are genes for which *s*(*E*) exhibits the largest *decreases* with increasing Ery concentration which are, therefore, likely due to media adaptation and not Ery resistance.

#### Polymorphisms under selection

*De novo* single nucleotide polymorphism (SNP) data observed significantly above 5% frequency at 120h indicate parallel, between-dosage adaptation (Supplementary §4, Figure 8). While a clinical case of *acr* evolution saw *de novo* protein structure SNPs in AcrAB^2^ we observed none of these in proteins that form the pump. Erythromycin binds to the ribosome and SNPs were observed in all treatments in ribosomal RNA operon *rrlC* but this also occurred in the absence of antibiotic suggesting media adaptation, not resistance. We found such putative media adaptations at all drug concentrations with SNPs in *fbaA* (a glycolytic enzyme), *glnK* (a nitrogen assimilation regulator), *acnA* (an oxidative stress protection and transcriptional regulator) alongside substantial polymorphism in phage genes *rzoD* (a lysis lipoprotein) and *ycbC* (a transmembrane protein) and *nohQ* (a putative packaging protein) which exhibit between-treatment parallelism (see Methods; for SNPs found in different biological replicates, see Figures S10, S11). Supplementary tables detail *de novo* SNPs above 5% in frequency at 120h that are unique to each antibiotic treatment, these indicate strongest selection for changes in amino acid transport and biosynthesis (genes *tauA* and *putP*).

**Figure 8.**
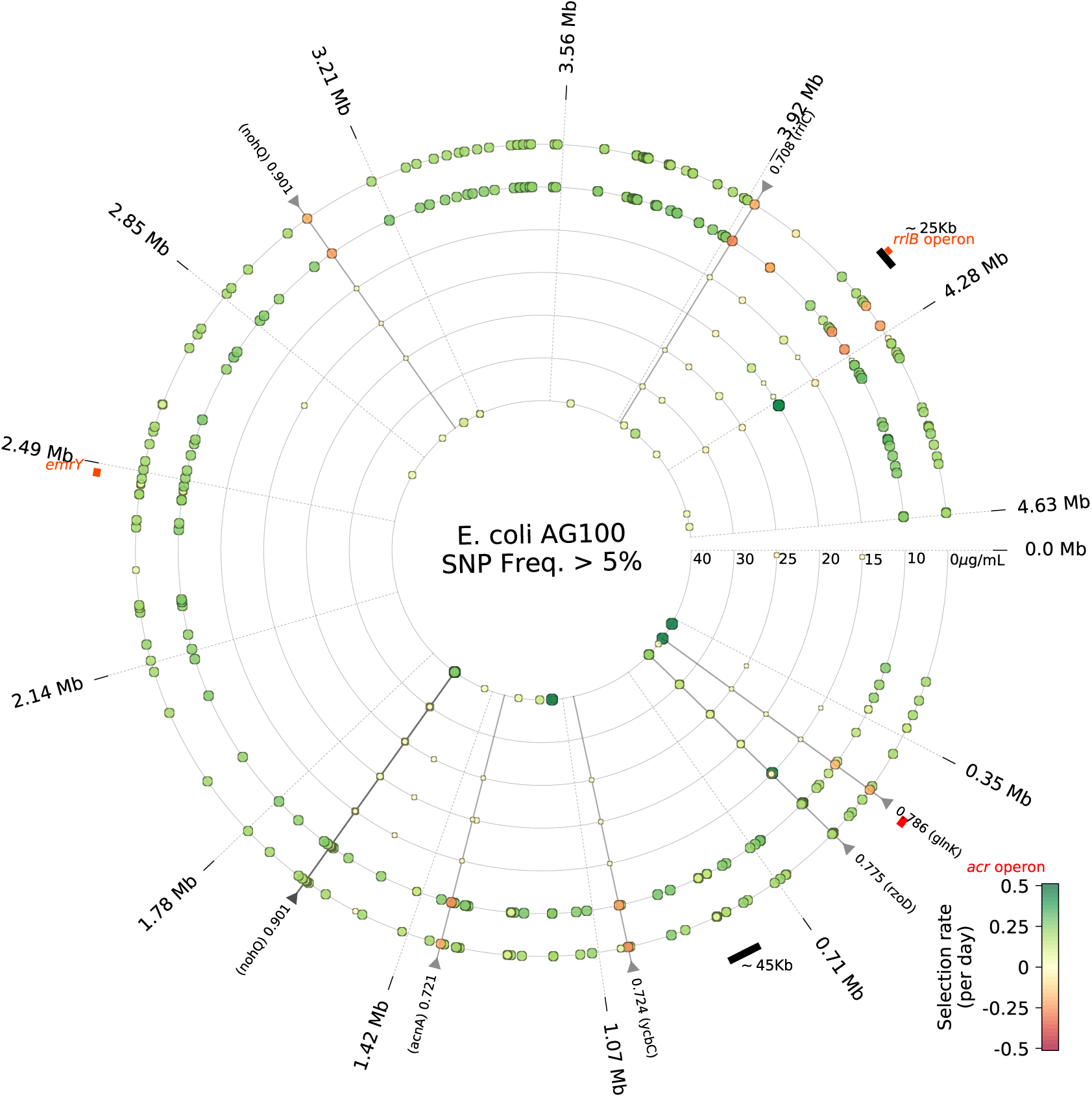
Whole genome sequencing data of AG100 at different erythromycin concentrations following 5 days’ exposure. *Single nucleotide polymorphisms*: Each ring highlights SNPs with dots whose colours indicate rates of sweep (see Methods). Triangles located in the perimeter indicate between-treatment parallelism coefficients based on day 5 frequencies (see Methods). Highly parallel evolution (SNPs with a parallelism coefficient above 0.7) is highlighted with a radial line. Regions of deletion or amplification are shown in the perimeter as black bands, the location of *acr* is a red band. Divergence between low and high dosage treatments is apparent from differences in the abundance of novel SNPs in each treatment (quantified in Figure S21).

We remark that it was not possible to use whole genome data to determine inverted-U geometries for SNPs associated only with resistance, as for this we would need parallel SNPs in all drug treatments that do not arise in the absence of antibiotic, but we found none.

#### The inverted-U is not ubiquitous as AG100A and TB108 do not exhibit one

We repeated the erythromycin treatment protocol using strain AG100A that has no functional AcrAB-TolC pump, finding it does not produce an inverted-U and, unexpectedly, exhibited most rapid population density at lower dosages (Figure S3). TB108^16^ has green fluorescent protein (GFP) fused to AcrB and, as its slightly reduced erythromycin MIC shows (Figure S1 shows TB108 has a reduced MIC compared with AG100 and eTB108), the AcrB-GFP fusion may reduce the efficacy of the pump. We found it too does not produce an inverted-U (Figure S12). We then adapted TB108 to 20*µg*/*ml* erythromycin until its MIC had increased to that of strain AG100 (Figure S1) and from here we isolated strain eTB108 which, upon applying the treatment protocol anew, did exhibit an inverted-U with its peak near the MIC dosage (Figure S13).

## Discussion

Microbial competition experiments have demonstrated that resistant strains can be selected from very low dosages upwards^5, 17^ and our data are consistent with this because we observe selection for resistance genes at all dosages assayed, low and high. The mutant selection window (MSW) hypothesis^18^ predicts selection for resistant subpopulations should be above at the MIC dose, but our data and theory do not agree and the MSW are marked in Figures 1 and 2 to illustrate this.

Indeed, we have shown that the dosage of most rapid selection for the amplification of regions of the genome implicated in antibiotic resistance can occur both below, and near, the wild-type MIC dose, depending on the function of the genes in question. This is consistent with both ours and prior theory^3^ which predict the peak of the inverted U could be found in principle at an arbitrary dose. Moreover, the theory of competitive release between sensitive and resistant subpopulations can explain^10^ the non-monotone dose responses we observe: the MRA occurs at the dose where susceptible sub-populations are suppressed, so they ‘release’ the maximum amount of extracelullar nutrients to resistant sub-populations at those dosages that do not suppress the growth of the latter. These theories indicate our findings are not specific to erythromycin and, indeed, prior *E.coli* data [19, Figure S13] shows an analogous dosing hotspot for the antibiotic doxycycline.

We have shown the inverted-U is not a universal property of resistance evolution and whether it exists, or not, exhibits subtleties. When an inverted-U is present, an *acr* knockout strain (AG100A) shows that its presence can be contingent upon a single gene. When the inverted-U is absent, our data show rates of adaptation can both increase and decrease monotonically with drug dose.

Increased *acr* expression has been associated with clinical resistance during sepsis^2^ whereby a two-fold increase in AcrB expression was detected after 1 week of antibiotic therapy. Our expression data exhibit similar timescales because a 3-fold amplification of *acr* here compares against the 2.5-fold change seen in that patient in one week. However, the large, 256-fold increase in MIC to erythromycin seen in the patient is not one we observe, indicating that other factors, such as cross-resistance resulting from multidrug therapy may play a role in hastening resistance in the clinic relative to this lab study. Antibiotic gradients within tissues^20^ is another possible feature of *in vivo* treatments that is absent from our model which could increase selection for resistance.

Our data show slower adaptation rates above the MIC, likely due to low population sizes, although the super-MIC recovery of some populations was observed (Figure S14 - S17). This leaves the question of how high must dose be so that no recovery is observed at all (*a.k.a* the mutant prevention concentration (MPC))? Accordingly, dosing erythromycin each day at 2xMIC produced no detectable growth (Figure S22) with an approximate 10^6^ cells per ml detection limit and thus, by treating above the MIC everywhere, at all times, we did prevent detectable growth. How relevant this observation is to clinical treatments is unclear, not least because spatiotemporal antibiotic gradients form in tissue^20^ where super-MIC dosages cannot be guaranteed.

We conclude that we have elucidated dosing hotspots and dose-dependent antibiotic resistance evolution using longitudinal data on large structural variants in the *E.coli* chromosome. However, as if to underline the difficulty of quantifying dose-dependent *de novo* evolution, we were unable to perform an analogous study for one, or more, single nucleotide variants by virtue of not being able to identify a SNP *a posteriori* that we could associate with a resistance phenotype, despite having phenotypic evidence of dose-dependent regulatory changes in the number of efflux pumps expressed per *acr* operon copy. Moreover, it was essential that evolved isolates were not cultured in the absence of antibiotic prior to DNA extraction, for had we done so, prior data quantifying the selection against *acr* amplifications^15^ indicates that the amplifications studied here would have been lost. Having designed our protocols to purposefully avoid that outcome, we have demonstrated how structural and single nucleotide variation can be mediated by an antibiotic to create dosages of most rapid resistance adaptation.

## Methods

### Strains and culture conditions

We used the strains of *Escherichia coli* AG100 (wildtype), AG100-A (*acrAB::Tn903*), both a gift from Stuart B. Levy, and TB108 (MG1655 *acrB-sfGFP-FRT*) grown in supplemented M9 minimal media. M9 was prepared by mixing dilute K_2_HPO_4_ (350g), KH_2_HPO_4_ (100g) in 1 L of de-ionized water and dilute of trisodium citrate (29.4g), (NH_4_)_2_SO_4_ (50g) and MgSO_4_ (10.45g) in 1L of de-ionized water, autoclaved and diluted accordingly. This media was supplemented with 0.2% (w/v) of glucose and 0.1% casamino acids from a filtered, sterilised 20% stock. All strains were incubated at 30*°*C and shaken linearly in a microtitre plate reader in 24h seasons. Optical culture density for all strains was read at 600nm and for TB108 and eTB108, GFP fluorescence was measured at 493nm/526nm (excitation/emission wavelenths).

### MIC assay and transfer protocol

A linear gradient of erythromycin was created by supplementing M9 media with different concentrations of erythromycin (Duchefa), using a prepared stock ranging from 0 to 50µg/mL. The cultures were incubated for 24h in 150µL of media with the corresponding drug concentration and transferred to a fresh plate after 24h of incubation using a 96-pin replicator (Nunc). Optical density and fluorescence readings were taking at regular 20 minute intervals and they were blank corrected by fitting the following three models to experimental data (raw data is shown in Figures S14–S17). If *B*(*t*) denotes the bacterial density at a given time, *t*, as measured by the microtitre plate reader, these models are

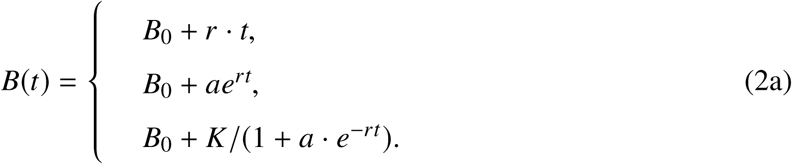

Here *B*_0_ denotes the summation of a blank and a population density reading at the beginning of the experiment following inoculation of the bacterium, *r* is the estimated *per capita* growth rate (in units of *h*^−1^, except for the linear growth model where it has units of bacteria per *h* and so is not measured *per capita*), *K* is the carrying capacity and *a* is a phenomenological coefficient. (Note that *B*(0) = *B*_0_ + *K*/(1 + *a*) which establishes that *a* is inversely related to inoculum density *B*(0) when *B*_0_ and *K* are known.) We implemented an algorithm in MATLAB that chose the best of these models (linear growth is appropriate in lag phase, exponential growth is appropriate for exponential phase, logistic growth is appropriate up to stationary phase) based on the lowest corrected Akaike Information Criterion (AICc). We then blank corrected the population density data using the corresponding *B*_0_ from the optimal datafit.

If *B*(*t*; *A*) denotes bacterial densities at time *t* cultured in the presence of a given concentration of antibiotic, *A*, we define the minimum inhibitory concentration (MIC) as the concentration satisfying *B*(24*h*, *A*) = *B*(24*h*, 0)/100, that is, the amount of drug required to inhibit 99% of the growth observed in the absence of antibiotic. These values are estimated (with error bars) by fitting a Hill function to dose response data (such as the fits shown in Figure S1 which provide examples of this).

### Whole genome sequencing

DNA extracted destructively from adapted populations (without any additional culture steps) was fragmented by sonication using a Biorupter for 30s on, 90s off, using low power for 10 minutes on ice. Libraries were prepared using SPRIworks cartridges for Illumina (Beckman Coulter) and Nextflex indexed adapters, with 300-600 bp size selection, amplified with 8 cycles PCR using Kapa HiFi DNA polymerase and purified using GeneRead kit (Qiagen). Concentrations were determined using a Bioanalyser 7500 DNA chip. Libraries were pooled in equimolar amounts, denatured, diluted to 6.5 pMol and clustered on a flowcell using a cBot (Illumina). 100 paired end sequencing with a custom barcode read was completed on a HiSeq 2500 using Truseq SBS v3 reagents (Illumina).

Reads were processed with fastq-mcf^21^ to remove adapters from the sequencing data and to trim and filter low-quality reads. Cycles with at least 1% of Ns were removed (command-line parameter: -x 0.01). The remaining reads were mapped to the AG100 reference genome using the Burrows-Wheeler aligner BWA^22, 23^ with standard parameters. The resulting alignments were processed with Samtools 1.3,^24^ with pair/trio calling enabled (command-line parameter: -T). Subsequently, alignments were sorted, artefacts and duplicates were removed, and finally the alignments were indexed. Unaligned reads were stored separately. Copy number variation was detected by analyzing coverage per base as measured by Bedtools^25^ after normalizing against the mean coverage. Finally, SNPs were detected using VarScan^26^ with the following parameters: p-value threshold of 0.05 for calling variants, minimum read depth of 20 to make a call at a position, base quality not less than 20 at a position to count a read, frequency to call homozygote of at least 0.9 (command-line parameters: `-p-value 0.05 -min-coverage 20 -min-avg-qual 20 -min-freq-for-hom 0.9`).

### Quantifying selection and between-dosage parallel genomic adaptation

We determined dN/dS data for all sequenced populations but did not find it helpful in terms of elucidating relationships between antibiotics dose and rates of adaptation.^27^ Thus, to quantify selection for genomic changes, we computed the rate of sweep by fitting the logistic function *<u>f</u>*(*t*) = 1/(1 + *p* · *e*^−*st*^) for *t* = 24*h*, 72*h* and 120*h* against longitudinal frequency data (noting that *f*(0) = 1/(1 + *p*) determines *p* and lim*_t_*_→∞_ *f* (*t*) = 1 for any *s* and *p*) and report the value of *s* as a proxy for the selection coefficient.

Between-treatment parallelism of a genomic change in gene g at a fixed timepoint is quantified by determining a vector ***f*** = (*f*_1_, *f*_2_, …, *f_N_*) of population frequencies at which that mutation is observed between *N* different treatments, we then determine the Euclidean distance, ***℘***, of ***f*** from a one-parameter family of uniform vectors of the form *λ* · (1, 1, …, 1), namely

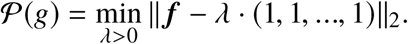

So, for example, if ***f***^(1)^ = (*f*_1_, *f*_2_, *f*_3_) = (0.49, 0.48, 0.47) for *de novo* SNP-1 in 3 treatments observed at 72h and ***f***^(2)^ = (*f*_1_, *f*_2_, *f*_3_) = (0.9, 0.8, 0.7) for SNP-2, we say that SNP-1 has a greater coefficient of parallelism between treatments because ***f***^(2)^ has more between-treatment variation than ***f***^(1)^ measured by the distance to the vector of perfect (i.e. zero) between-treatment variation, namely *λ* · (1, 1, 1) for some scalar *λ*.

We also determine 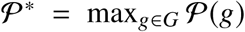 where *G* is chosen using a selection criterion for genomic regions of interest and we then define the coefficient of parallelism, *p*(*g*), relative to 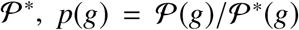. Here we use a value of *p*(*g*) > 0.7 to highlight the most highly parallel mutations where *G* is chosen to reflect only those mutants for which each *f_j_* > 0.05. Such thresholding criteria are arbitrary and were merely chosen so that Figure 3B would be as legible and informative as possible.

To produce Figure 7, we fitted the above logistic function to raw Illumina coverage data for each gene in the *E.coli* genome to quantify selection coefficients (in units of change of relative coverage [i.e. relative to the mean across the genome] per day - this is called the *selection proxy* in Figure 6) on changes in gene copy number at each dosage. If this selection coefficient, which depends on erythromycin dosage, is labelled *s*(*E*) for erythromycin, E = 0, 5, 10, …, 40*µg*/*ml*, we then applied quadratic regression to *s*(*E*) data to model it as a quadratic function and so to predict the dosage at which selection was maximal. We plotted one datum per gene for which the maximal selection rate was significantly positive to produce Figure 7.

### Evolving TB108 to recover AcrB function

The MIC of the GFP-tagged strain TB108 was observed to lie between the MICs of AG100 and AG100-A (Figure S1), suggesting a possible reduction in function of the pump AcrAB-TolC likely due to GFP being physically fused to AcrB. To recover the function of the pump we therefore propagated a culture of TB108 exposed to 10µg/mL of erythromycin and subcultured for 5 days, tracking changes in MIC. We considered the complex GFP-AcrAB-TolC to be functional when the MIC of the evolved strain, a new strain deemed eTB108, was restored to a value close to that of the wildtype AG100. Figure S1 shows eTB108 and AG100 have similar dose responses to erythromycin, both with significantly higher MICs than TB108. Figure S18 then uses fluoresence micropscopy image data made of single cells sampled from an adapted population to corroborate the spectrophotometer / plate reader data using a different measurement technique that eTB108 is capable of doubling its per-cell GFP-AcrB expression level when propagated in the presence of erythromycin. We therefore used eTB108 in our study to quantify the dynamics of adaptation in GFP-AcrB expression.

### Oscillations in fluorescence data

Oscillations in AcrB-GFP expression data were observed in Figure 2E on the timescale of an hour, or so, so we sought the mechanism supporting these within our protocols. To quantify the frequency of these oscillations we computed the discrete Fourier transform of the relative fluorescence dataset from Figure 2E using the `fft` function implemented in MATLAB 2014a. As a result, two dominant sub-harmonics were observed, one with a frequency of ∼10h and the other of ∼0.75h (Figure S19). In negative control cultures that were not inoculated with bacteria we only observed the latter. The former harmonic, of lower frequency, which corresponds to the relatively slow timescale up-regulation of *acr* that takes place in exponential phase whereas the latter harmonic, of higher frequency and observed in the absence of biological material, must therefore be an electrical or mechanical phenomenon induced into the liquid media by the microtitre plate reading devices.

### Quantifying rates of adaptation

Our definition of rate of adaptation (ROA) is taken from the literature:^12^ the ROA is a rate of change estimate of any dynamic phenotype measured for a bacterial population, for instance population GFP expression levels or growth rate. By way of example, suppose *r*(0) denotes the *per capita* growth rate at the first measurement of a longitudinal bacterial culture experiment and suppose *r*(*t*) is growth rate at time *t* > 0. We define the adaptive time, *t_a_*, as the time that satisfies *r*(*t_a_*) = *r*(0) + Δ*r*/2 where Δ*r* = *r*(*T*) − *r*(0) (where *T* is the time at the end of the experiment) and then the rate of adaptation is *α* := (Δ*r*/2)/*t_a_*. The rate of adaptation data we present (e.g. Figures 2A-B) is the result of plotting α at different drug concentrations. We also compute α where the timeseries, *r*(*t*), here is replaced with data on GFP per OD which we use as a proxy for AcrB expressed per cell (e.g. Figure 2C).

We used two methods to quantify *per capita* growth rate to verify the robustness of our statements about rates of adaptation and their dependence on drug dose. The first method is based on a the so-called forward difference approximation (*r_e_*) of derivatives that we apply to the best model fit, *B*(*t*), obtained using (2a) applied to bacterial population density data. The second method uses a reciprocal area under the curve (*r_auc_*) measure which has units of *h*^−1^ to reflect a per capita growth rate. These values are defined as

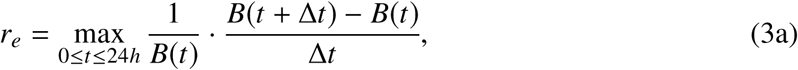

where Δ*t* = 20mins because bacterial densities are read with that frequency, and

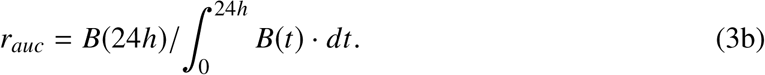

Rates of adaptation computed using *r_e_* are shown in Figure S13 whereas all other figures in the main text use *r_auc_*. Note that both *r_e_* and *r_auc_*can be applied directly to data if *B*(*t*) is a raw longitudinal population density dataset, but applying the model fit first, as we do here, provides more robust estimates of these statistics by effectively using the model to filter high-frequency noise. This is important as growth rate is a derivative of *per capita* population density and any noise present in the latter will be exacerbated under the operation of taking derivatives.

### The effective antibiotic dose (EAD)

The EAD provides a way of quantifying a temporal reduction in drug efficacy without having to measure MIC changes of all adapted populations, something that would be prohibitively time consuming to undertake due to the number of treatments involved in this study (each treatment round for each replicate induces a potential change in MIC each day). The question that defines the EAD is this: isolate an adapted bacterium that has been treated after *n* rounds of treatment with antibiotic at dosage *A*_∗_ and which is observed to grow to some population density, *B*_∗_ at that time; how much antibiotic would have been required so that, when applied to the wild-type bacterium on treatment round 1, it too would have grown to *B*_∗_ units? Given that a dose-response, *B*(*A*) which describes population density as a function of antibiotic dose over 24h, is *a priori* known for the wild-type, this verbal description is asking that we find the value *A_EAD_* for which *B*(*A_EAD_*) = *B*_∗_.

When *B*(·) is a monotone decreasing function represented theoretically by a Hill function, then *A_EAD_* = *B*^−1^(*B*_∗_) and the aforementioned monotonicity property implies that *B*^−1^ (the inverse function of *B*) is monotone decreasing too. As *B*(*A*) is a decreasing function of dose *A* and if *B*_∗_ > *B*(*A*_∗_) because of resistance adaptation, it follows that *A_EAD_* = *B*^−1^(*B*_∗_) < *B*^−1^ *B*(*A*_∗_) = *A*_∗_, so that *A_EAD_* will be smaller than *A*_∗_; this is the effective reduction in dose. In the text we measure the value of this EAD following five rounds of treatment where *A*_∗_ is taken to be the MIC of the wild-type bacterium (Figures 1D and S4).

### Data availability

Whole-genome data is published and publicly available on ENA (https://www.ebi.ac.uk/ena) with accession number PRJEB28068. All other data will be made publicly available on acceptance of the paper. All raw data from evolutionary assays are shown in supplementary figures.

## 3 Supplementary Figures

**Figure S1.**
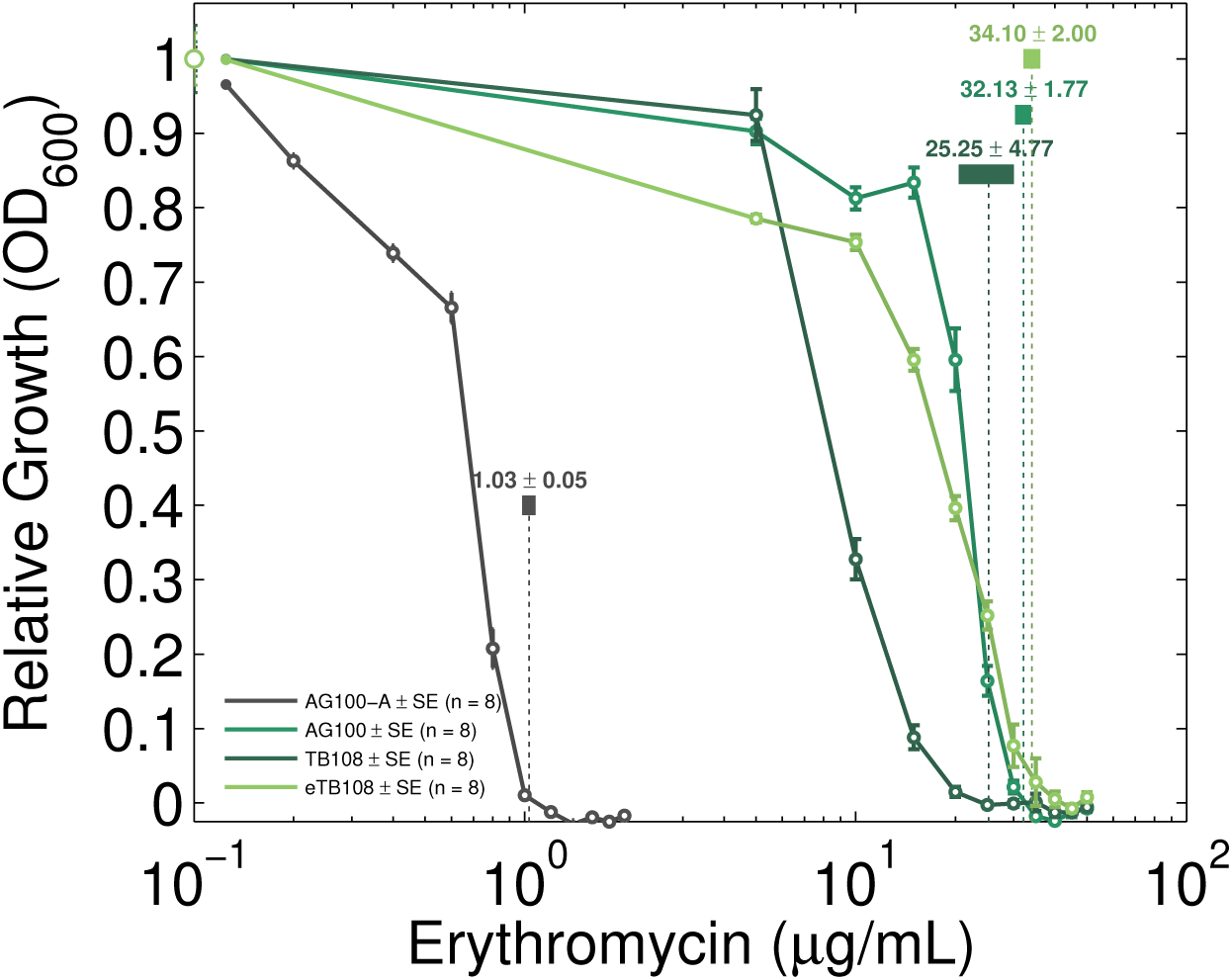
Dose-response profiles for all the strains used. Four erythromycin dose-response profiles for the strains of *E. coli* AG100, AG100-A, TB108, and eTB108 where optical density data has been measured at 600nm (OD_600_) after 24h of growth. OD_600_ is shown on the y-axis whilst the concentration of erythromycin is represented in a logarithmic scale on the x-axis. The IC_99_ and its 95% confidence intervals, determined using n = 8, are shown for each strain and indicated as horizontal bars.

**Figure S2.**
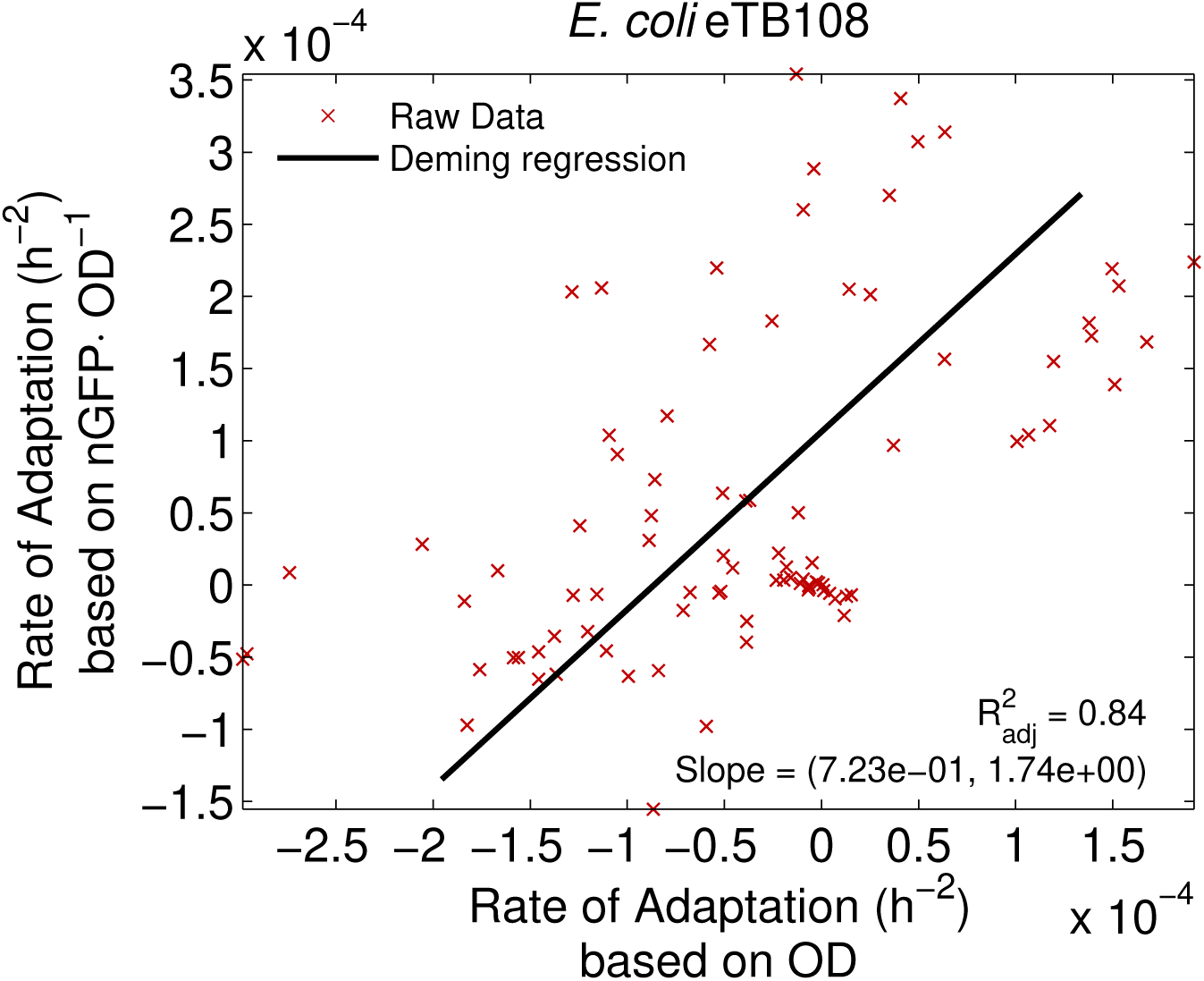
The union of the entire adaptation dataset of strain eTB108 collated at all drug dosages shows that the rate of adaptation of population density (OD) correlates positively and significantly with the rate of adaptation of AcrB-GFP expression (Deming regression, linear slope parameter 95% CI ≈ (0.72, 1.74)).

**Figure S3.**
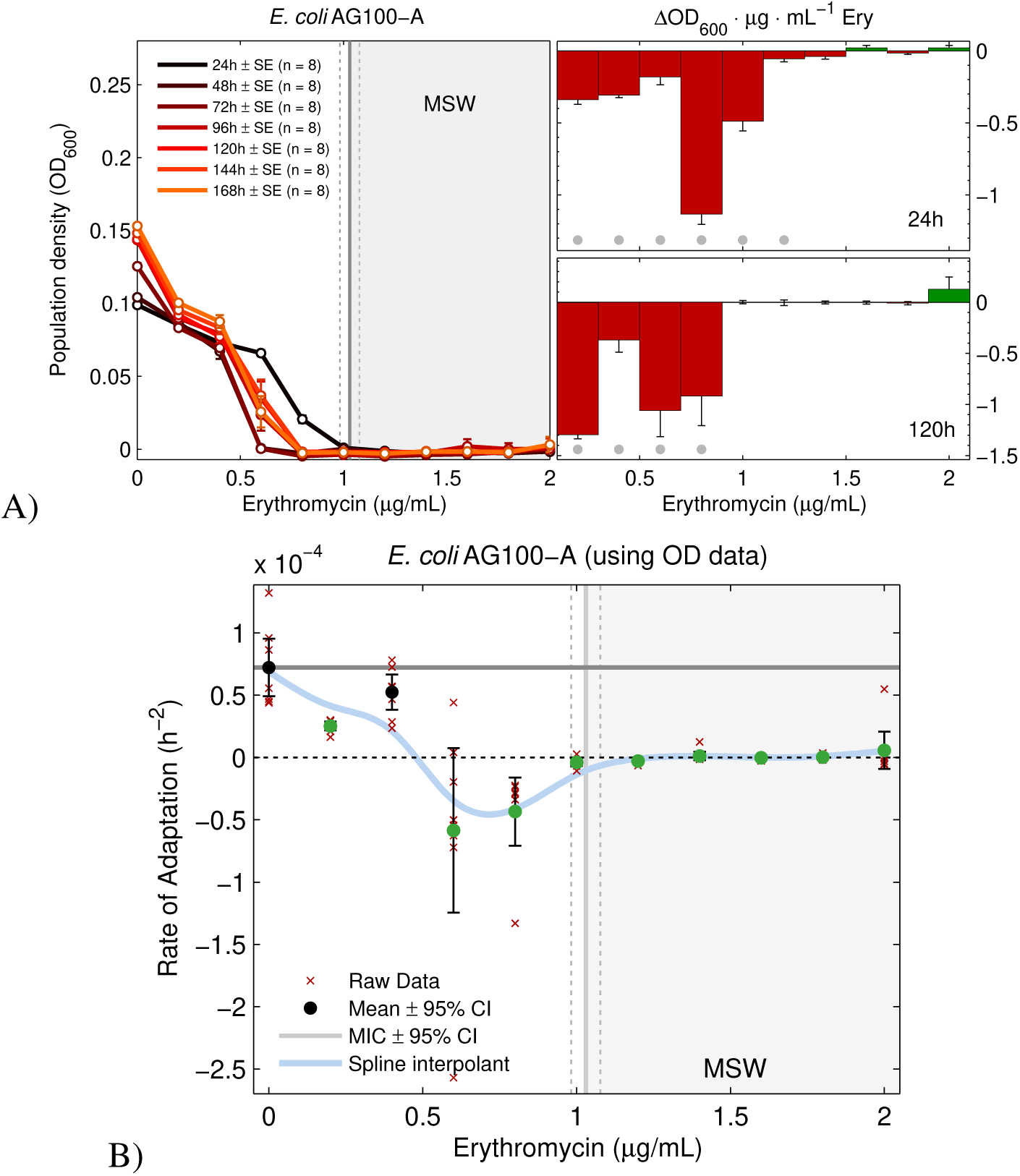
Evolving the dose response of erythromycin and *E. coli* AG100A. **A)** The leftmost subplot illustrates daily dose-response profile data. The MIC and MSW, with a 95% confidence interval and measured after 24h as per standardised sensitivity tests, is represented by dark grey and dashed lines. The putative MSW is shown as a light grey area. The effect on bacterial density of increments in drug concentration are shown in the two rightmost subplots after 24h and 120h of exposure to erythromycin (mean ± standard error, n = 8). Significant changes, with respect to the null hypothesis of *no change induced by increasing drug concentration* (from a 2-sided t-test) are denoted by a grey dot. **B)** Rates of adaptation at different drug concentrations determined using *r_auc_*. The dashed line is the point no detectable adaptation, the thick, continuous grey line highlights the adaptation to the media in the absence of erythromycin. Significant changes with respect to this datum are denoted by green dots based on two-sided t-tests (mean ± standard error, n = 8).

**Figure S4.**
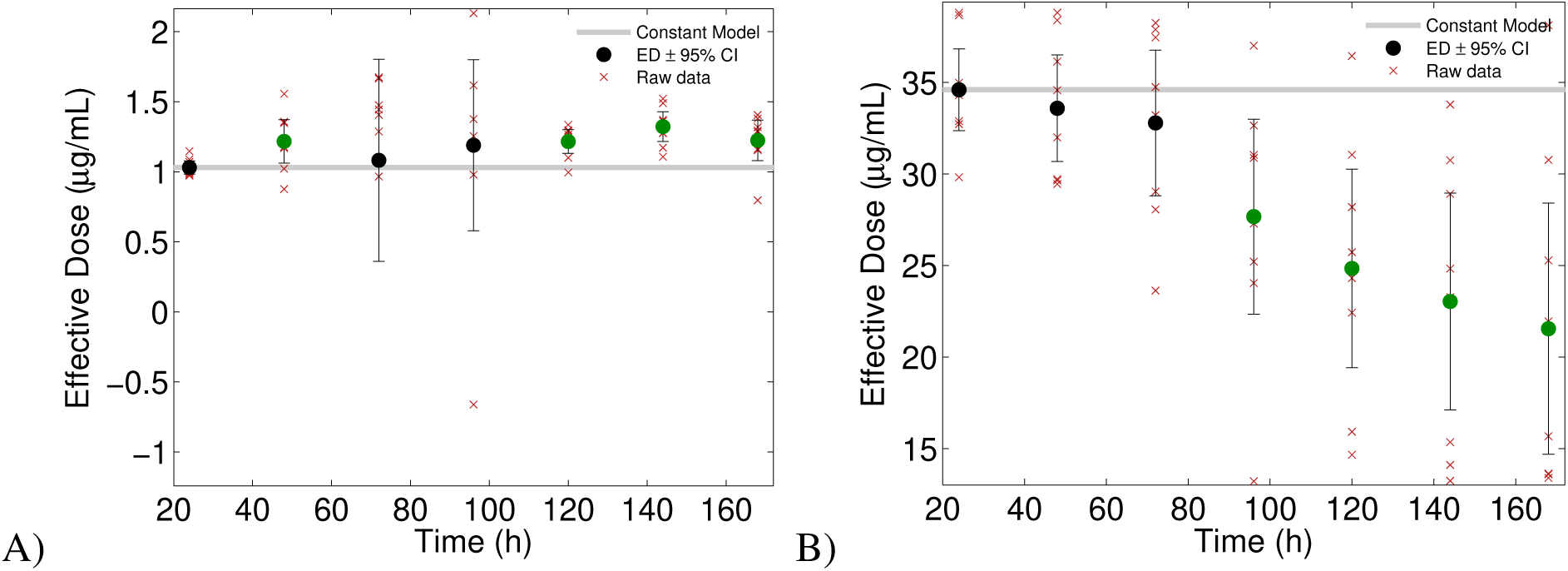
Dynamics of the effective antibiotic dose (EAD). A) The EAD does not change for the AG100A strain when this is propagated in the treatment protocol described in the main text and methods. B) The EAD does change when eTB108 is treated in this way, reducing in value by approximately 50%. Significant changes based on a 2-sided t-test with *p* < 0.05 are shown as a green dot, non-significant changes are shown as a black dot.

**Figure S5.**
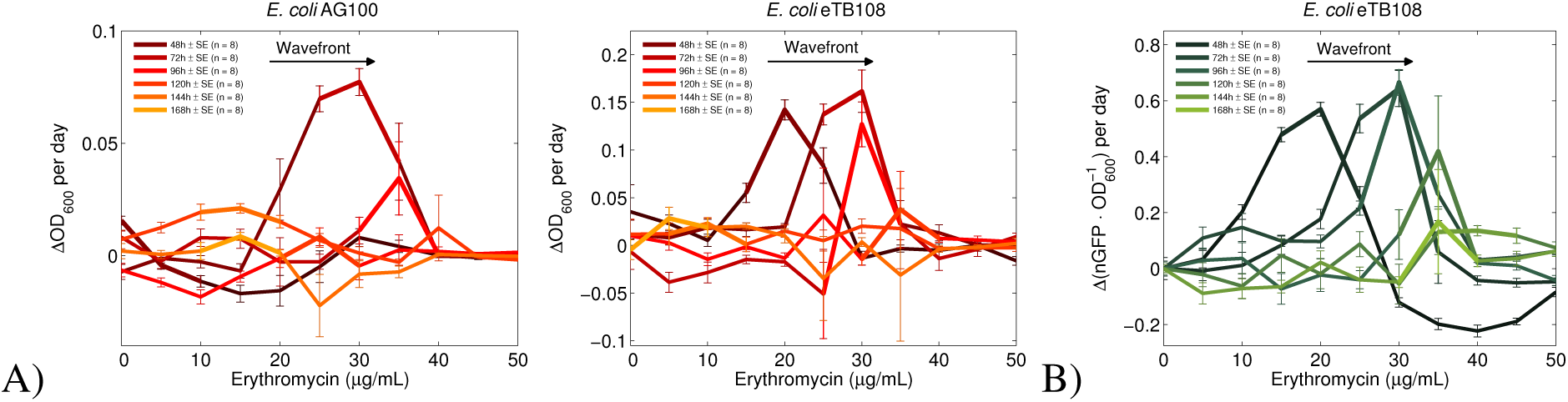
Daily changes in bacterial density at different drug concentrations for 3 strains. Beginning on day two, each line in A and B represents the difference in bacterial density (orange) or relative GFP fluorescence (green) between a given day and the preceding day’s data. For strain eTB108, comparing data in A and B indicate a potential correlation between the adaptation profile of bacterial density and efflux pumps per cell, where a proxy for the latter in B is the daily change in GFP units per OD signal. This correlation is quantified in Figures S13 and S2.

**Figure S6.**
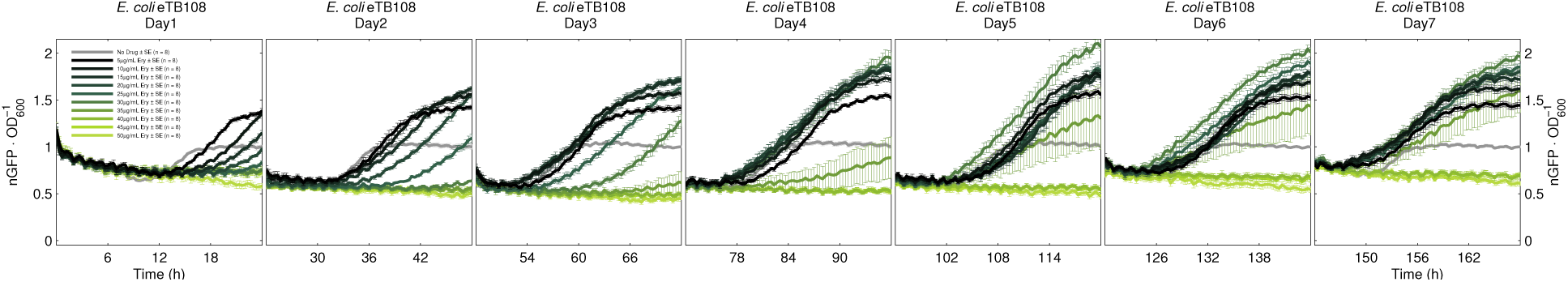
The changing dynamics of GFP per OD (for strain eTB108) shown here are a proxy for mean AcrB-GFP per cell dynamics, shown for 7 daily treatments and showing within-season dynamics from lag to stationary phases at all of the antibiotic dosages used. Note the logistic-type shape of each curve and the increasing AcrB-GFP expression through time each day. By the end of the experiment, only the very-highest dosage treatments have not seen a rise in mean AcrB-GFP levels. Note also the grey curve, this is the antibiotic-free control line and the AcrB-GFP expression pattern remains the same each day, also forming a logistic-like curve.

**Figure S7.**
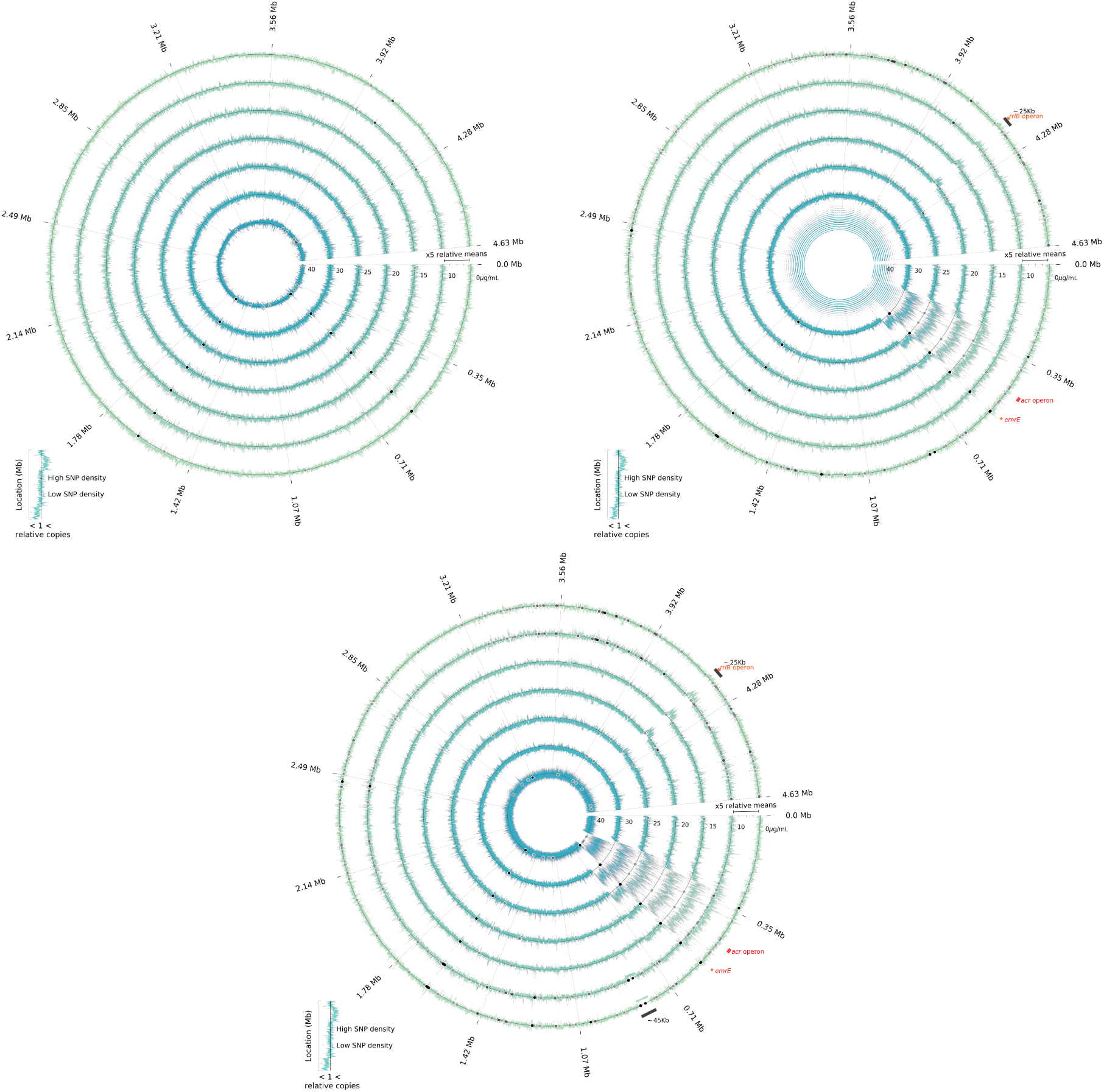
Temporal variation in coverage data at different erythromycin concentrations (Replicate 1) Normalised Illumina coverage data of AG100 metagenomes exposed to different erythromycin concentrations after 1 (top-left), 3 (top-right) and 5 (middle bottom) daily treatments. The outmost ring shows data for cells propagated in the absence of antibiotic, followed by rings showing data for increasing concentrations of drug (10-40µg/mL); Figures S8 and S9 show analogous data for two more replicates. Larger vales (visible spikes) represent genomic amplification, smaller values (visible dips) indicate gene deletions. Single nucleotide polymorphism (SNPs) are indicated by dots in the corresponding chromosomal location. The operon *acr*, responsible for the multidrug efflux pump AcrAB-TolC, is highlighted in red between 480,553-485,707 bp. Other signatures, like the loss of ∼45Kb (804,268-_3_8_6_49,700 bp) and the amplification of a ∼25kb region between 4,164,724-4,189,435 bp, are highlighted in black. The former region contains putative phage-like genes whereas the latter is rich in metabolic genes, including the ribosomal operon *rrlB*, 50S ribosomal protein L1 (*rplA*) and the RNA polymerase subunit β (*rpoB*).

**Figure S8.**
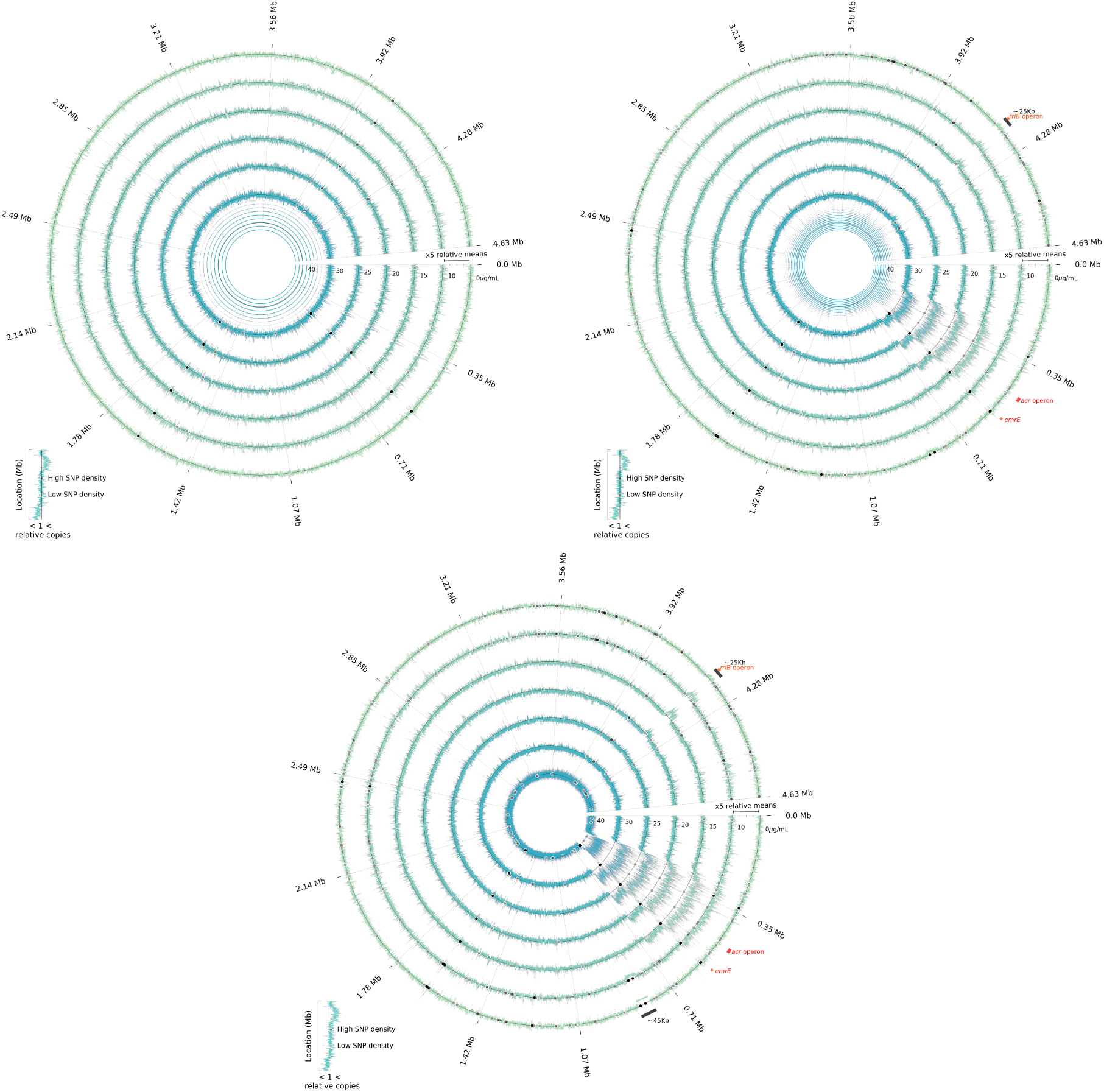
A technical replicate of Figure S7.

**Figure S9.**
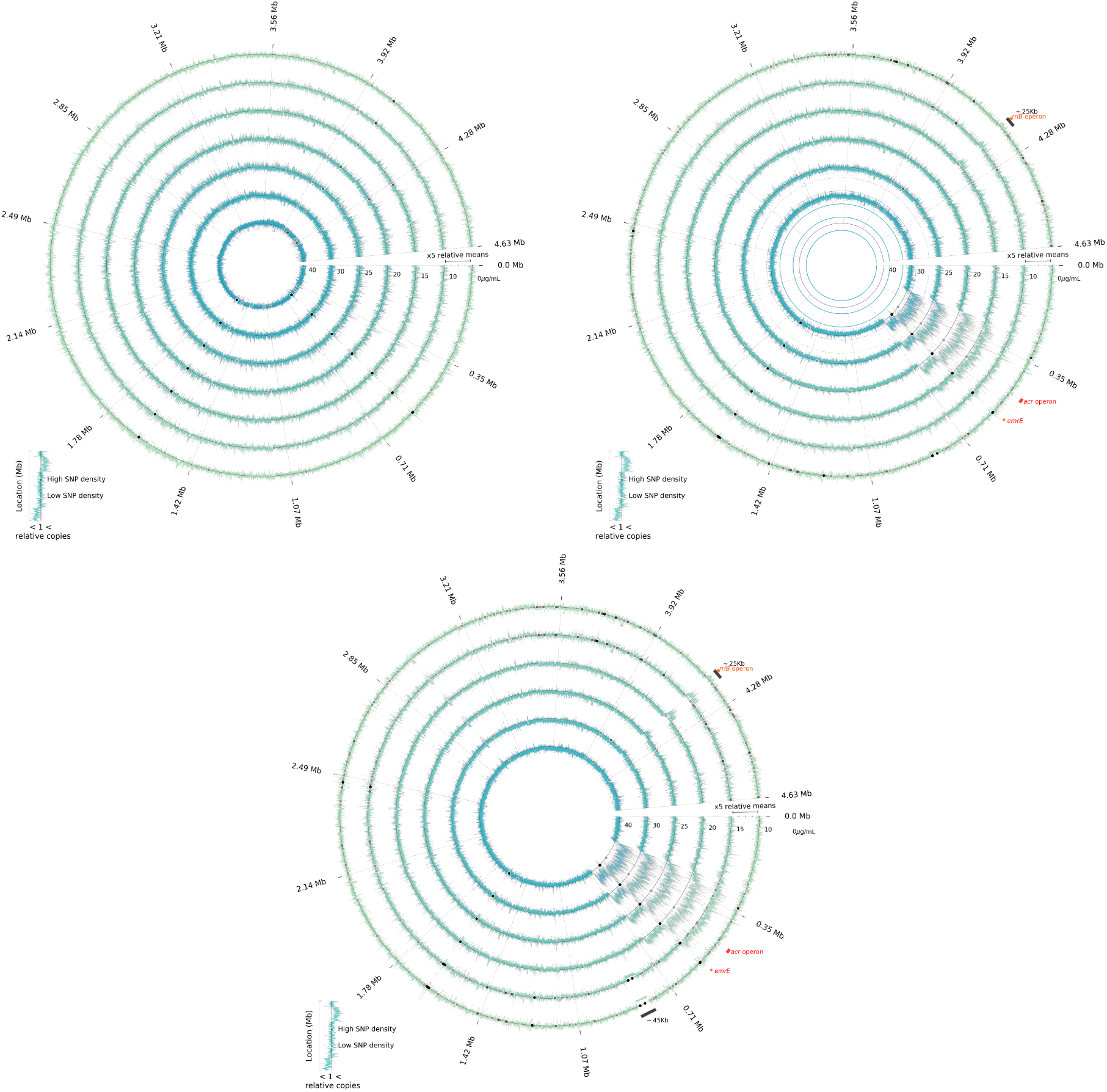
A technical replicate of Figure S7.

**Figure S10.**
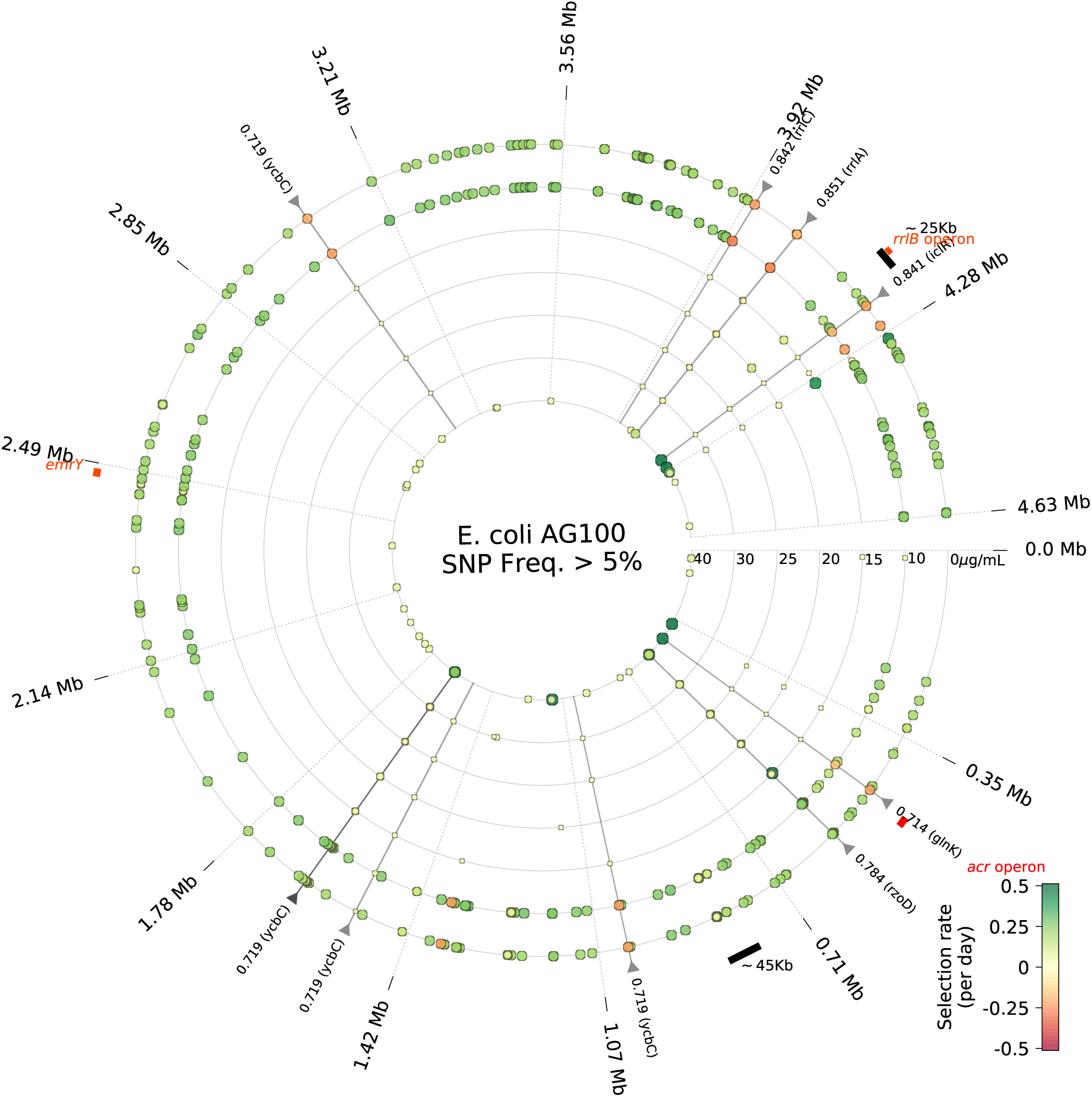
A technical replicate of Figure 3.

**Figure S11.**
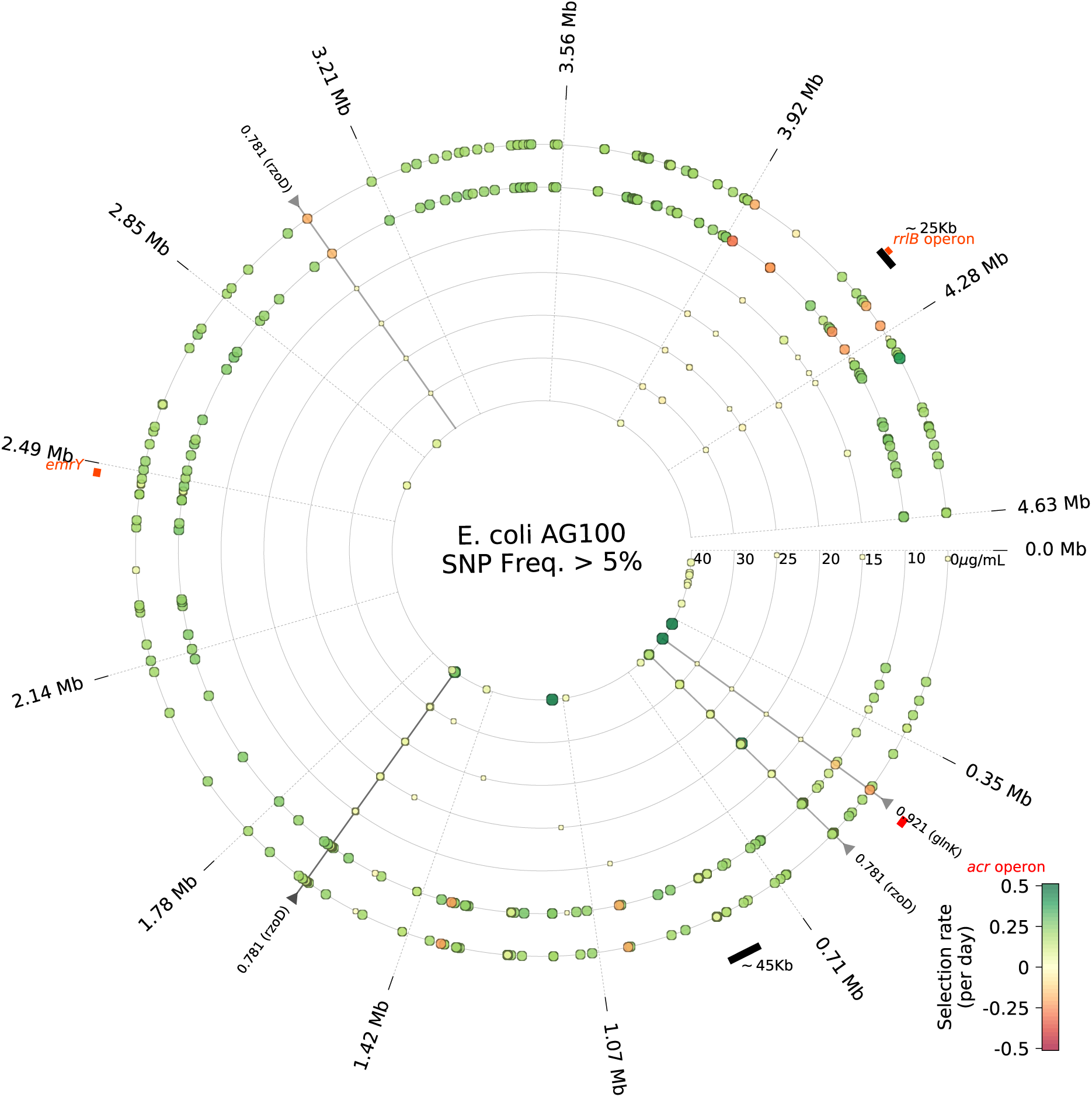
A technical replicate of Figure 3

**Figure S12.**
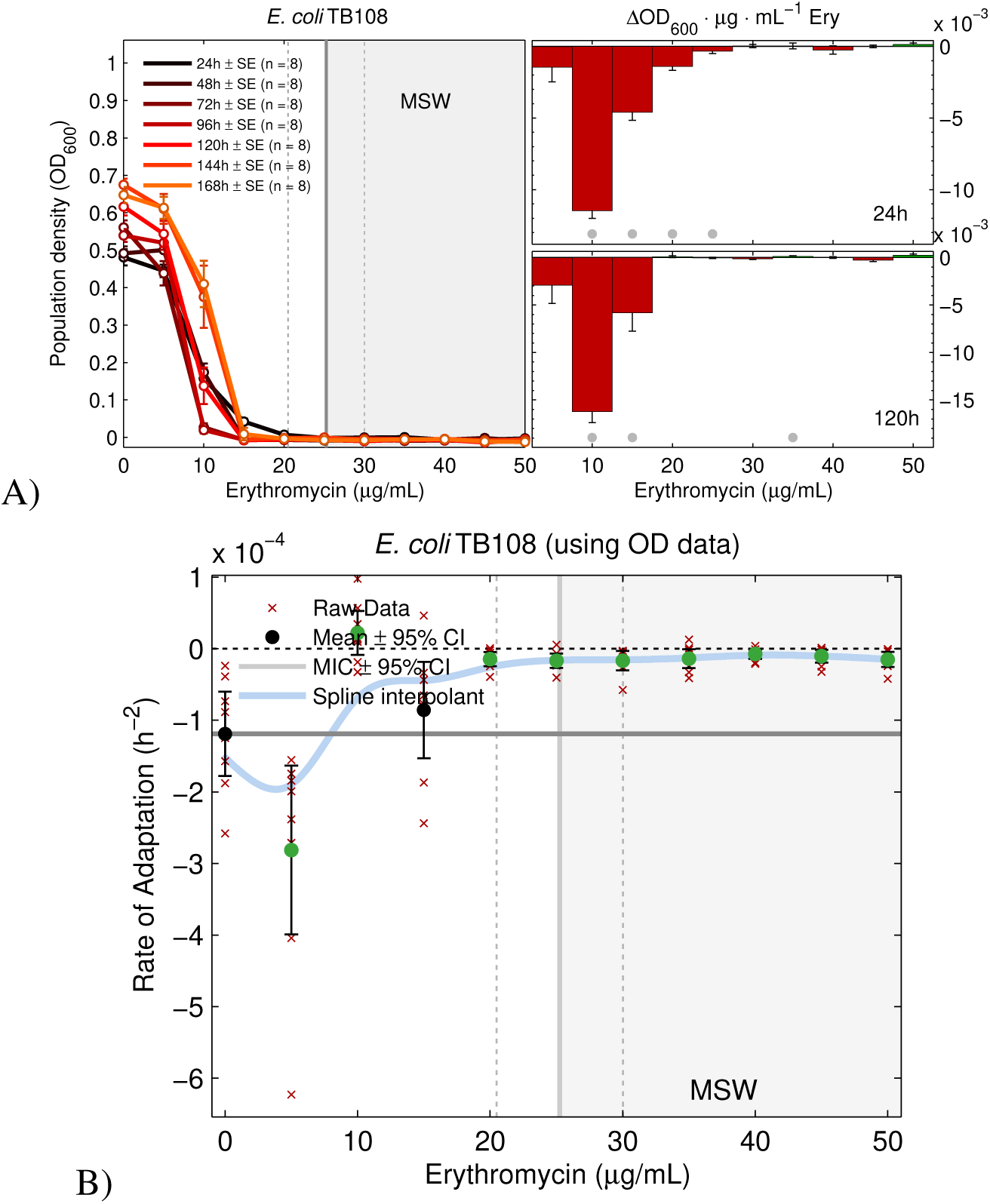
Evolution in response to erythromycin of *E. coli* TB108. A) Erythromycin dose-response profiles for *E. coli* TB108 measured every 24h. The growth measured as OD_600_ is represented in the y-axis as a function of the concentration of erythromycin, represented in the x-axis. For the subplots, the y-axis represent the point-to-point slope changes of the dose-response profiles and significantly positive (green) or negative (red) slopes are highlighted accordingly (using *p* < 0.01). Here, all such data is negative and thus increasing dosages decrease population densities at all times. B) The rate of adaptation statistics determined from data in A shows an increasing form whereby population density adaptation increases with drug dose.

**Figure S13.**
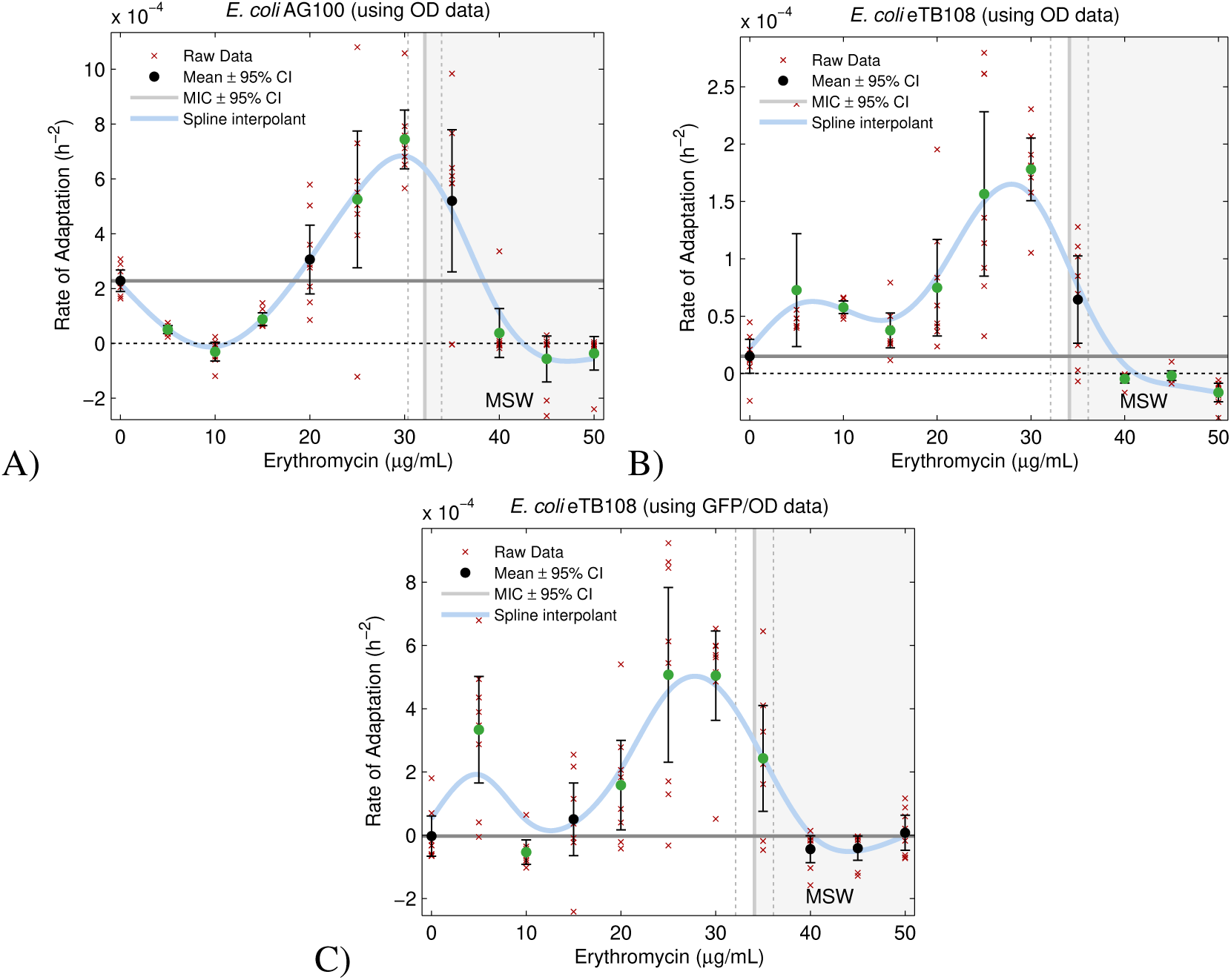
Rates of population density adaptation using an alternative metric (*r_e_*). Rates of adaptation determined using *r_e_* (see Methods) which shows the variation in rates of adaptation with drug concentration. A and B use cell density (OD) data whereas C uses the mean relative abundance of AcrAB-TolC protein (GFP per OD). The light grey line is a baseline determined by propagation in media without antibiotic whereas the thick, continuous light blue line shows adaptation to media with erythromycin. Significant differences with respect to the baseline are noted by green dots based on two-sided t-tests (mean ± standard error with n = 8).

**Figure S14.**
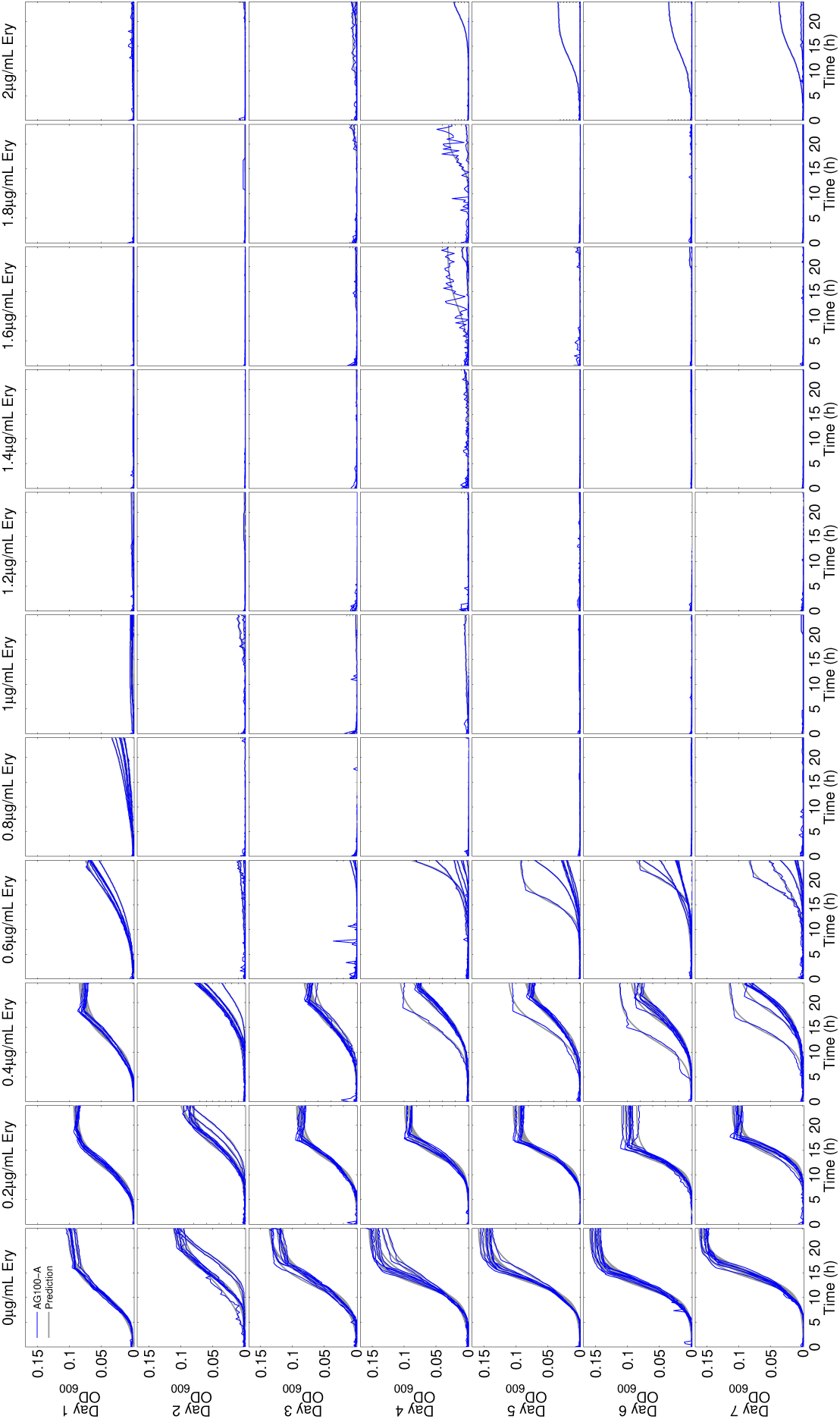
Raw optical density data and logistic model fits for all treatments of AG100A. Growth profiles for *E. coli* AG100A based on optical density measured at 600nm (OD_600_) every 20min, for 24h, for 7 days. Each subplot contains the growth profile of eight replicates through time, presented as a function of the dose of erythromycin (column-wise). Different rows show the model fit (blue) and the empirical data (grey) for different days ranging from 1 to 7.

**Figure S15.**
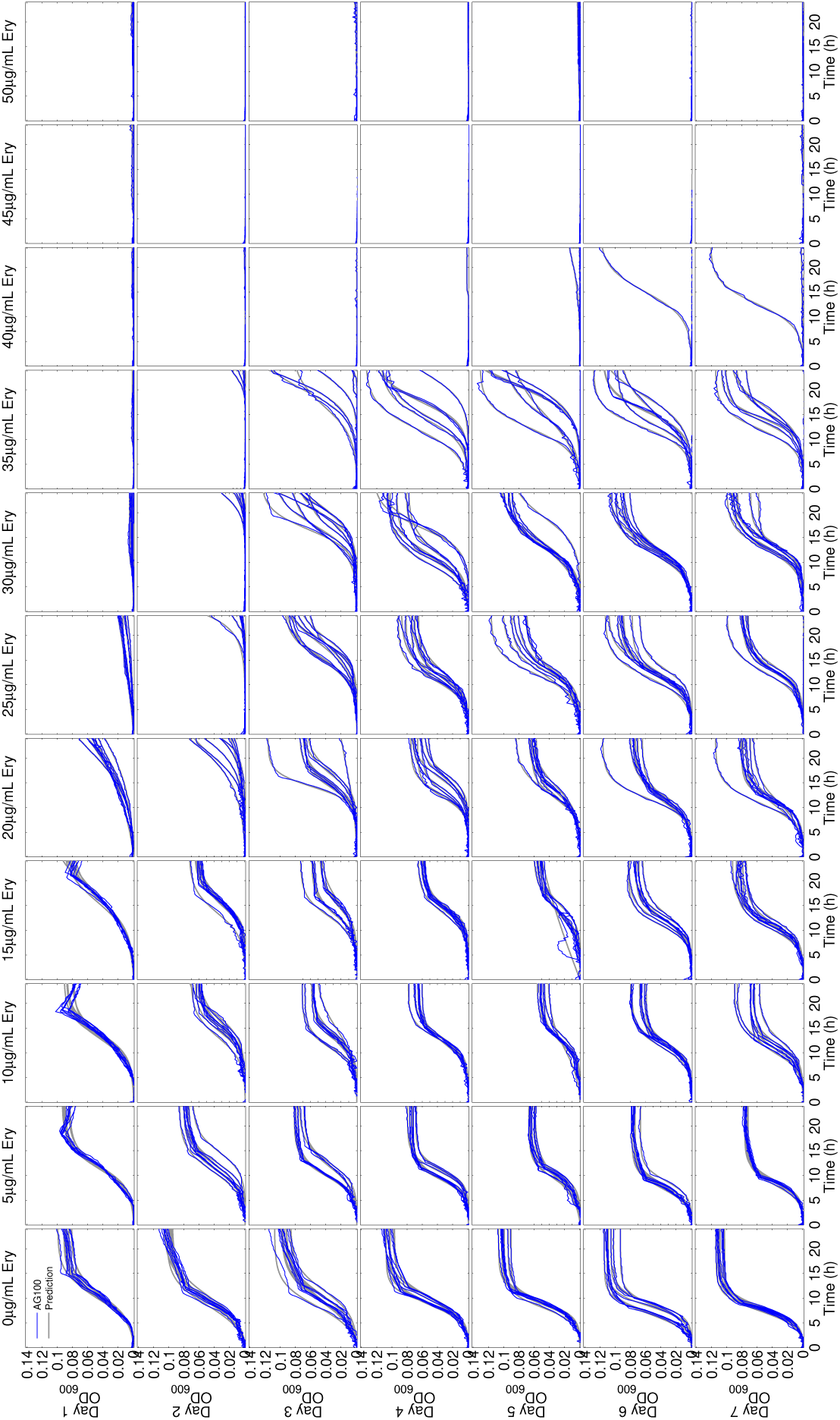
Raw optical density data and model fit for all treatments of AG100. This is analogous data to Figure S14 but for the strain AG100. Note the MIC is around 30µg/*ml* of erythromycin but significant population growth is observed in some replicates up to 40µg/*ml*.

**Figure S16.**
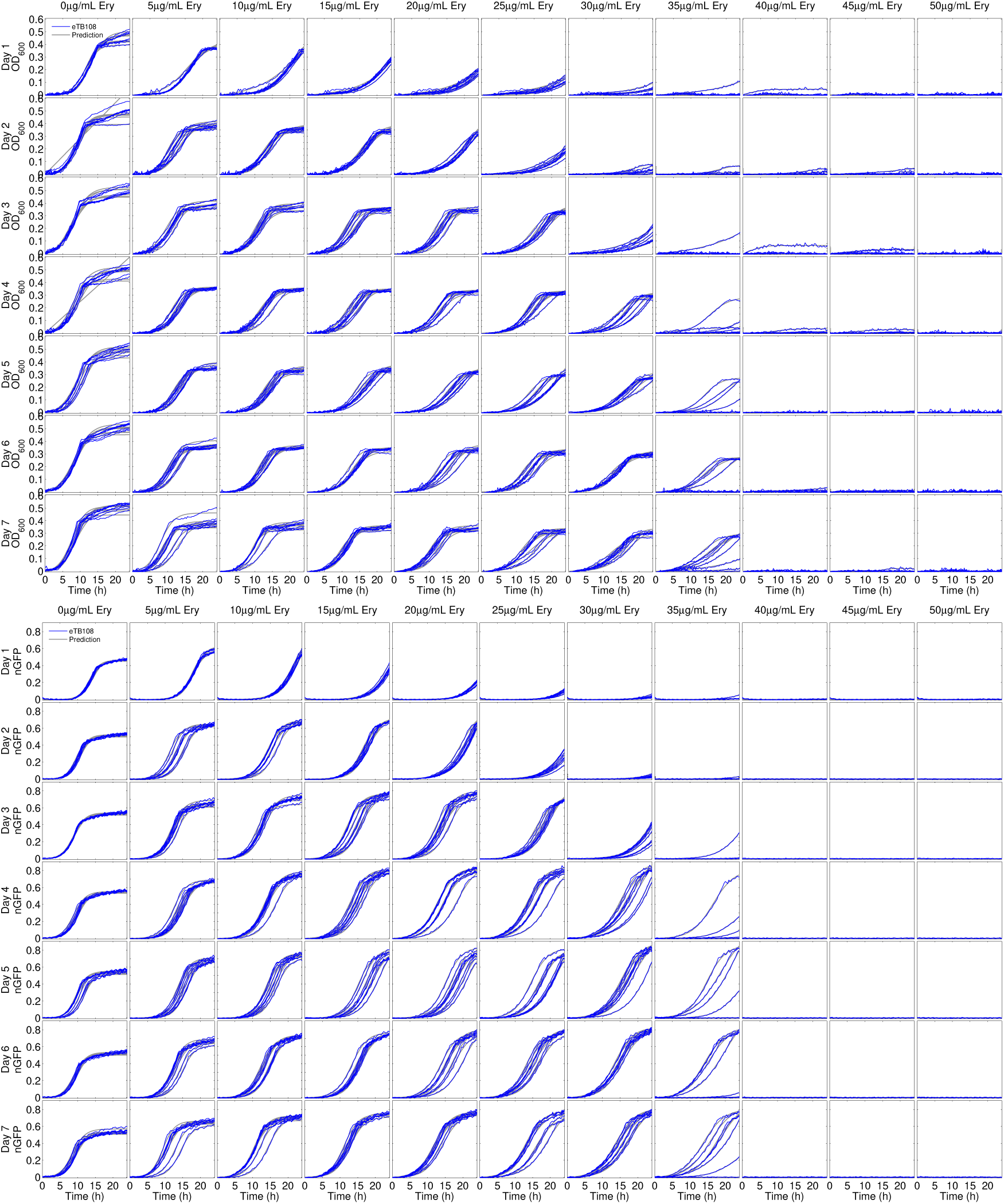
Raw optical density (top) and fluorescence (bottom) data and associated model fits for eTB108. Growth profiles are shown for *E. coli* eTB108 based on optical density measured at 600nm (OD_600_, top), and normalised fluorescence (nGFP, bottom), each taken every 20min for 7 days. Subplots contain data profiles of eight replicates as a function of time shown as a function of erythromycin column-wise. Different rows show fit and data for different days, from 1 to 7.

**Figure S17.**
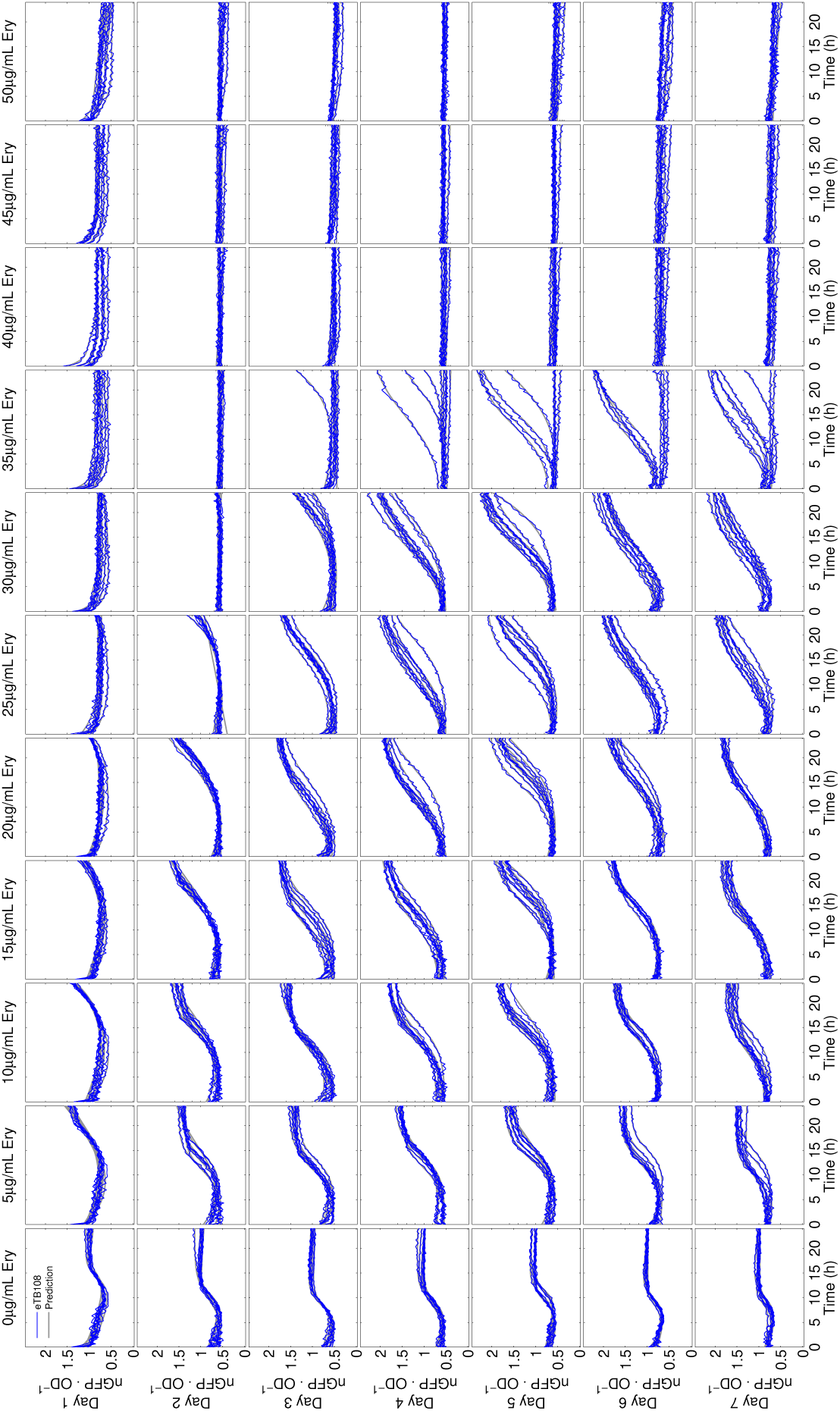
Raw relative fluorescence data and associated model fits for eTB108. Growth profiles for *E. coli* eTB108 based on relative fluorescence per OD_600_ unit (nGFP · OD^−1^) read every 20min for 7 days. Each subplot contains the growth profile of eight replicates as a function of time, shown as a function of erythromycin column-wise. Different rows show the model fit (blue) and data (grey) for days 1 to 7.

**Figure S18.**
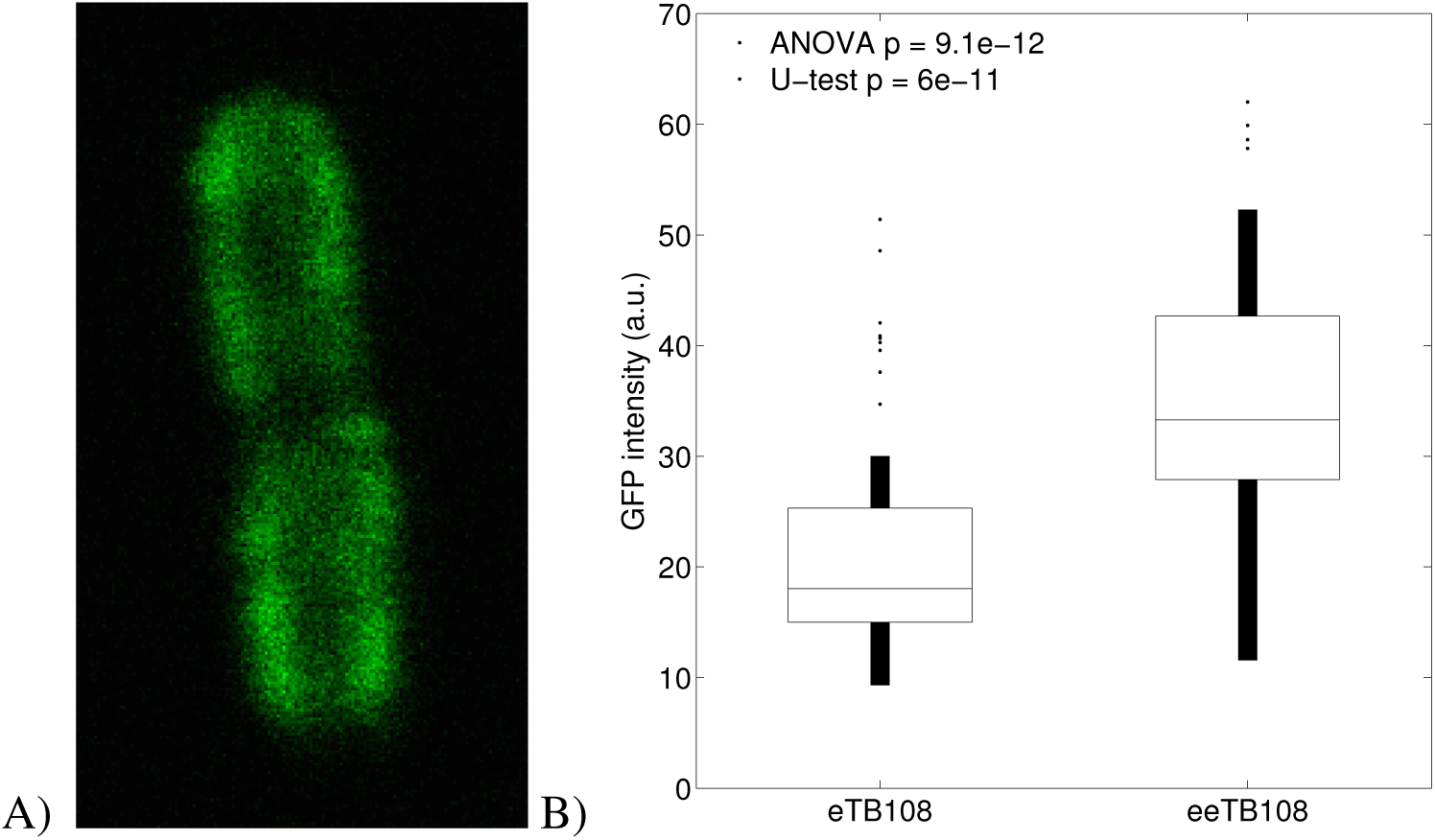
A demonstration that 2-fold increases in AcrB expression occur during therapy using microscopy. A) Fluorescence microscopy demonstrates the GFP-fused AcrB protein localises in the outer membrane of *E.coli*. B) Fluorescence microscope image data taken before and after erythromycin treatment show eTB108 is capable of doubling the expression of AcrB-GFP when propagated at sub-MIC levels (20*µg*/*ml*) for 5 days, thus producing a strain that we denote eeTB108 for the purposes of this figure. The GFP levels reported here are measured per cell using a fluorescence microscope that are quantified using segmentation algorithms in Matlab.

**Figure S19.**
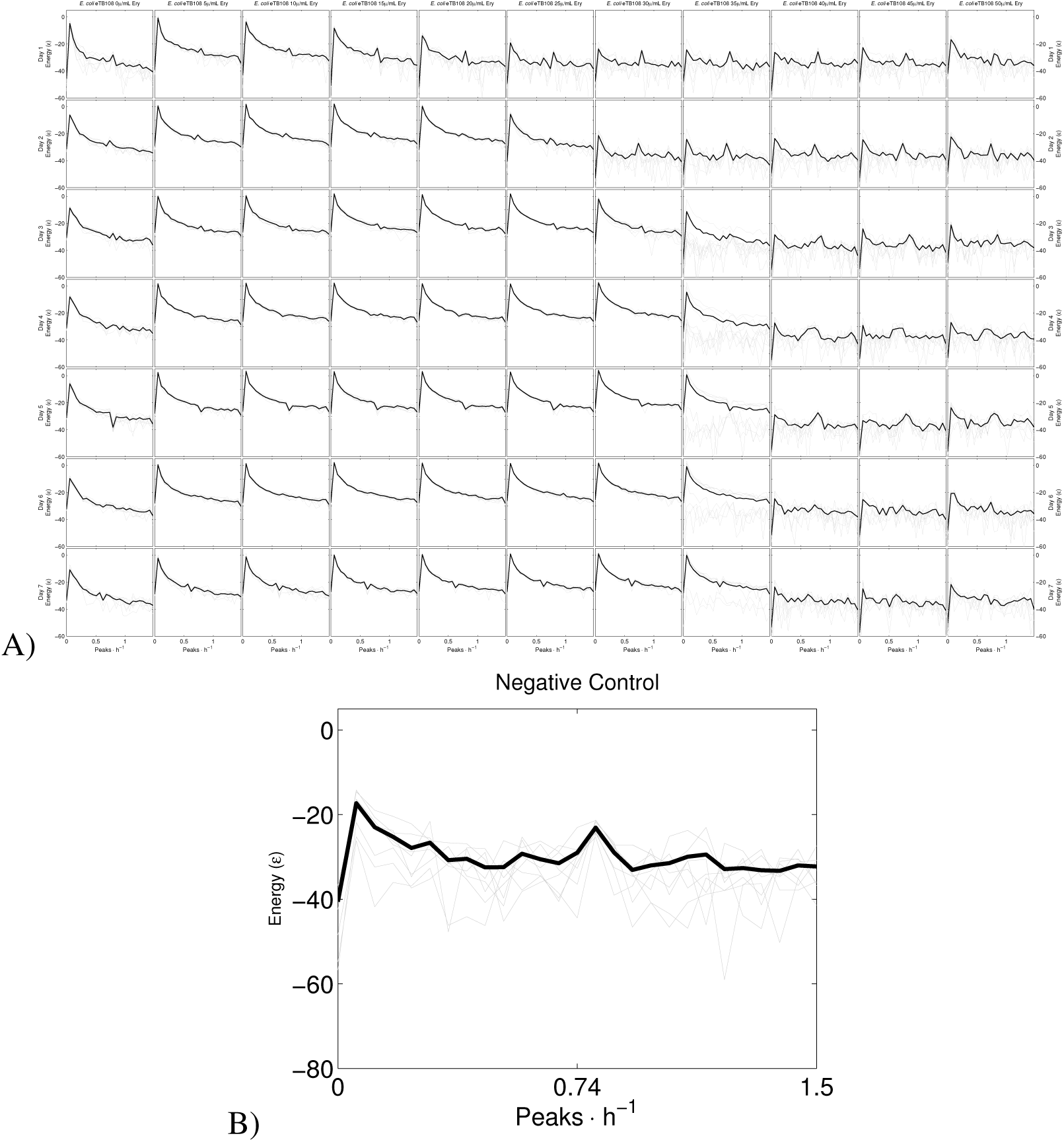
Fast Fourier transform (FFT) spectra for eTB108 GFP per OD data. **A)** The frequency of oscillations is shown on the x-axis and the energy at each frequency on the y-axis. Columns represent different days, different concentrations of erythromycin are shown in rows. Two features are common here: a peak every ∼3/4 h and another every ∼10 h. The former feature corresponds to the small oscillations observed in Figure 2D. **B)** The same information but for the non-inoculated microtitre plates (i.e. the negative control) also show a peak near 3/4h. We conclude that oscillations in GFP per OD data (e.g. Figure S6) do not have a biological basis but are a (presumably) mechanical or electrical feature of the spectrophotometer device that produces these datasets.

**Figure S20.**
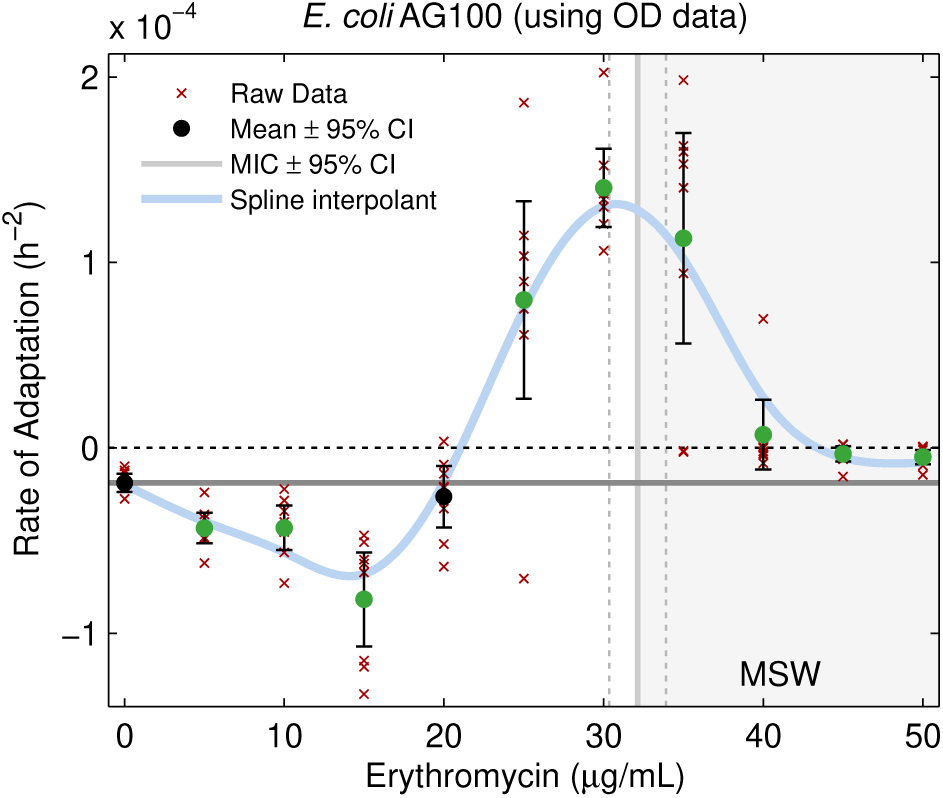
Rates of population density adaptation depend on drug concentration. Rates of adaptation for population density of strain AG100 (measured using *r_auc_* applied to OD data) vary with drug concentration, peaking near the MIC.

**Figure S21.**
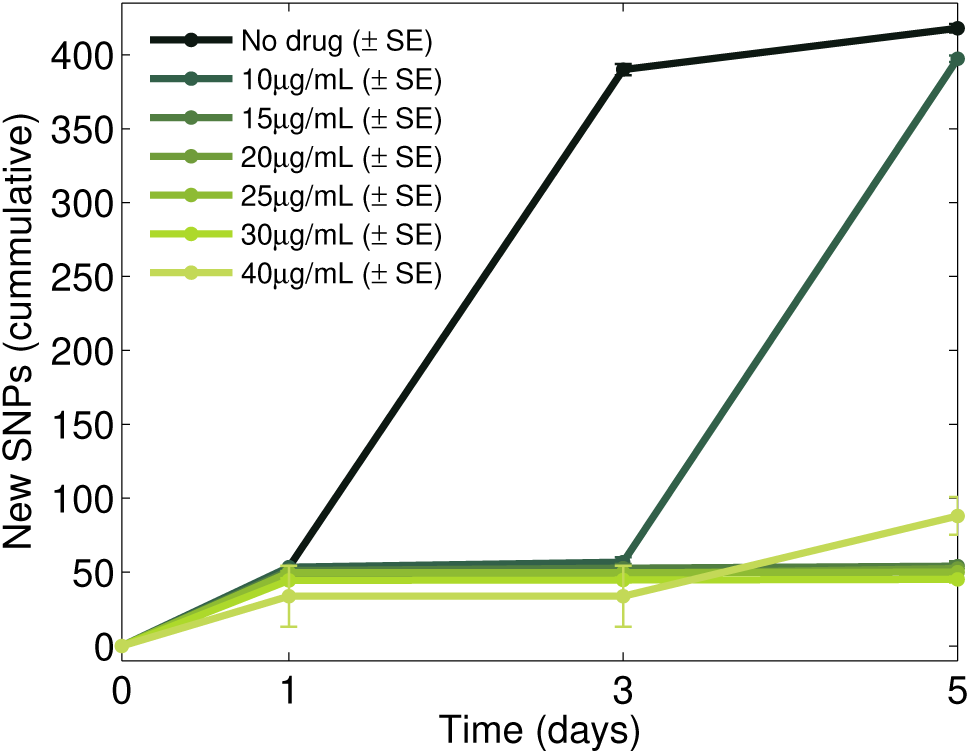
Cumulative novel SNPs detected per genome in each treatment on days 1, 3 and 5. Most novel SNPs are observed in the absence of antibiotic where population densities are highest, so higher drug dosages lead to fewer novel SNPs each day.

**Figure S22.**
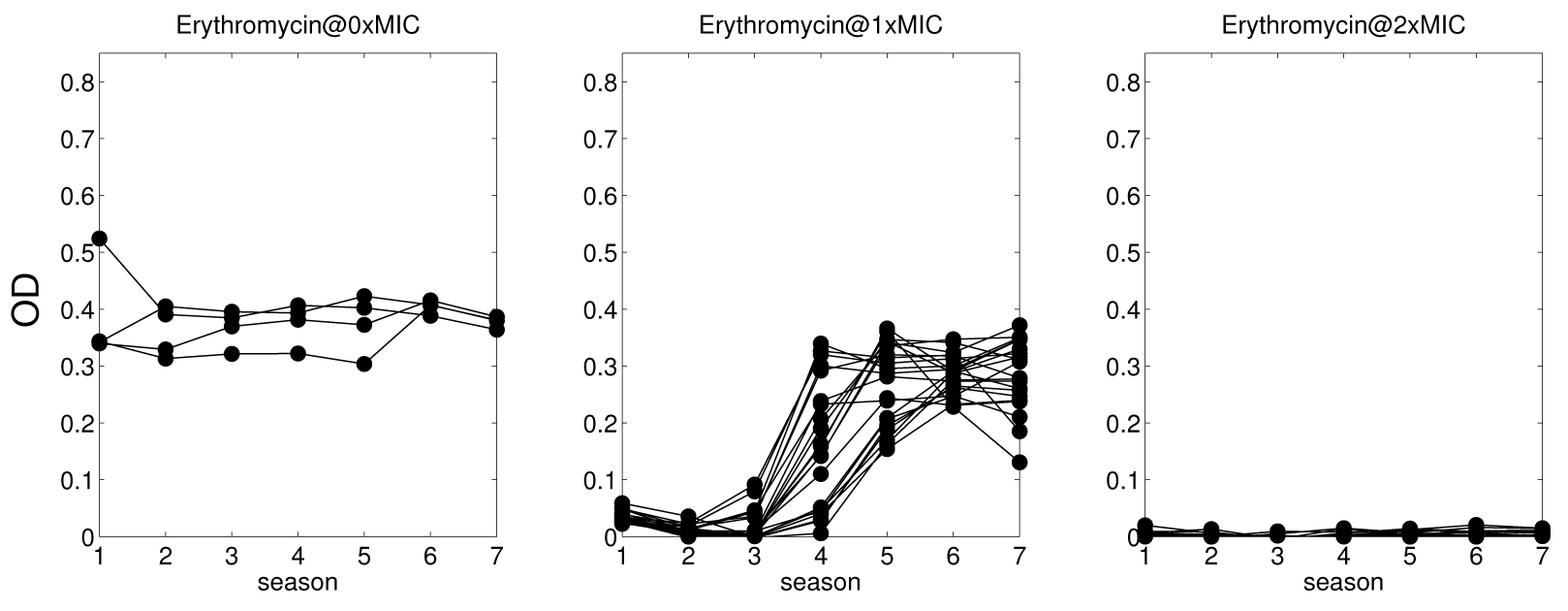
No growth for a high dose erythromycin treatment. AG100 optical density (OD) data for 7 × 24h seasons of erythromycin treatment are shown, implemented at 0xMIC (drug free), 1xMIC and 2xMIC dosages, respectively. In the last of these we observe no significant (above background noise floor) growth detected in a microtitre plate reader (16 replicates).

## 4 Supplementary Tables

The following 6 tables show novel SNPs detected (i.e. not present in the wild-type genome at inoculation) that are above 5% frequency in their respective populations sampled from each of the different antibiotic treatments that are not also present in any of the drug-free control treatment replicates. The value of *s_j_* in each case is a selection coefficient determined from the frequency dynamics of each detected SNP, where the *s*-coefficient is the rate of fixation of that SNP determined from a fit of a logistic model to that frequency data in each case.

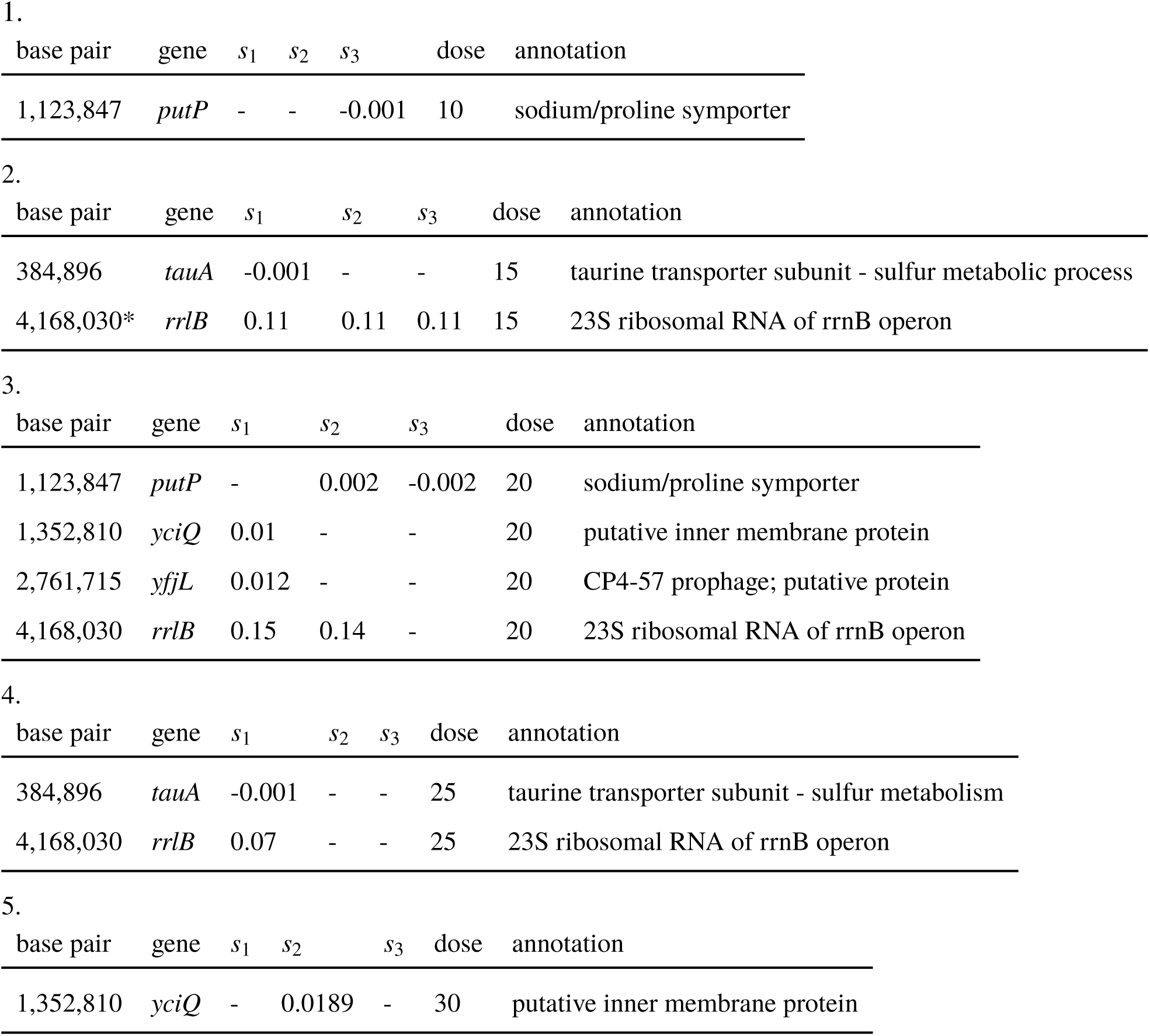
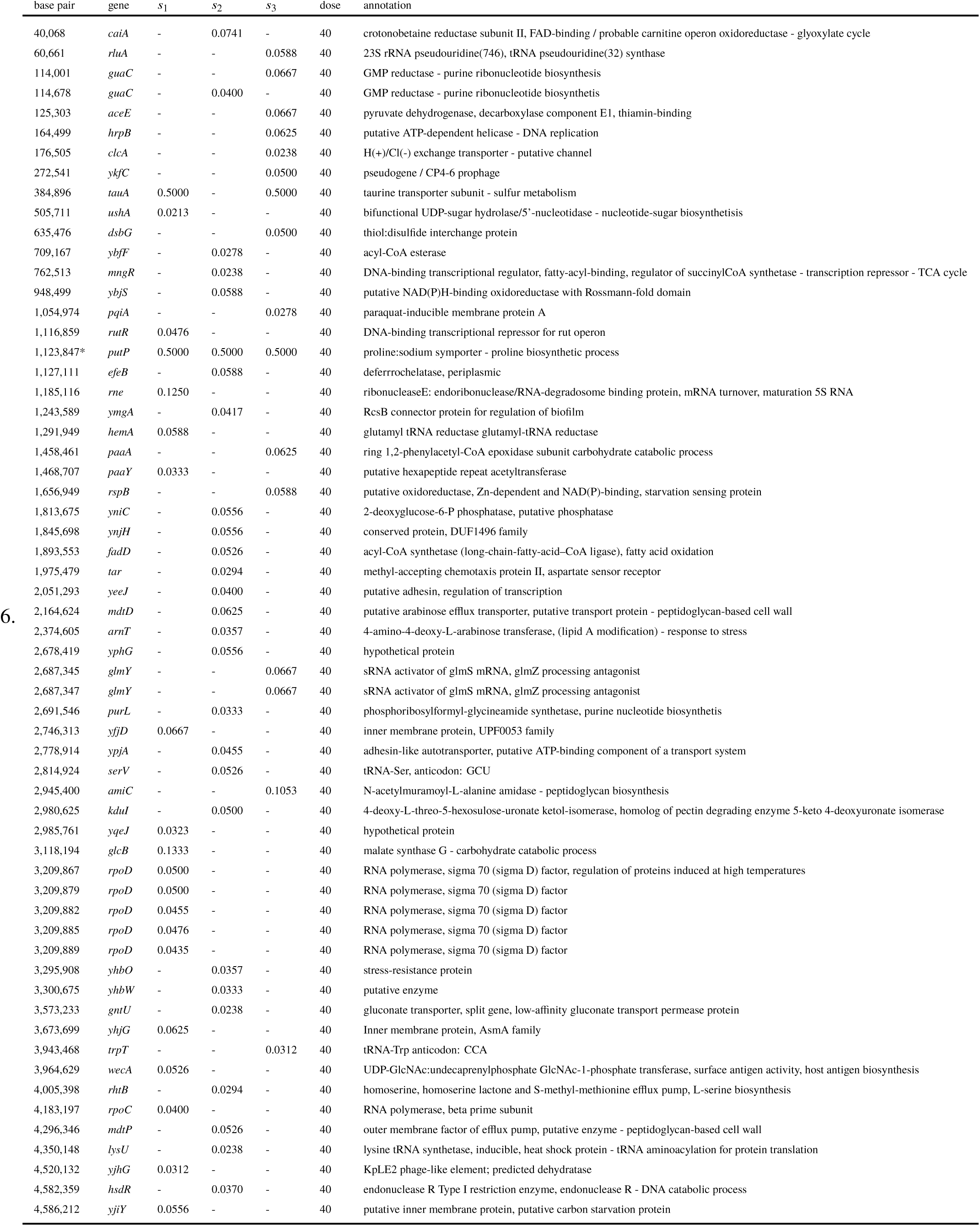

